# The Dachsous/Fat/Four-jointed pathway implements axial long-range cell orientation

**DOI:** 10.1101/092650

**Authors:** Federica Mangione, Enrique Martín-Blanco

**Affiliations:** Instituto de Biología Molecular de Barcelona, Consejo Superior de Investigaciones Científicas Parc Científic de Barcelona, Baldiri Reixac 10-12, 08028 Barcelona, Spain

**Author notes:** Corresponding author: Enrique Martin-Blanco.

## Abstract

Despite a cumulative body of knowledge describing short-range cell interactions in morphogenetic processes, relatively little is known on the mechanism involved in the long-range spatial and temporal coordination of cells to build functional and structurally organized tissues. In particular, the attainment of a functionally optimized epithelia must require directional cues to instruct cell movements and cell orientations throughout the tissue field. In *Drosophila,* the adult epidermis of the abdominal segments is created *de novo* by the replacement of obsolete larval epidermal cells (LECs) by histoblasts (imaginal founder cells). As these proliferate, expand and fuse, they uniformly organize orienting on the surface along the antero-posterior axis. We found that the coordinated, axially oriented changes in shape of histoblasts respond to a dynamic, yet stereotyped redesign of the epithelial field mediated by the Dachsous/Fat/Four-jointed (Ds/Ft/Fj) pathway. The establishment and refinement of the expression gradients of the atypical cadherins Ds and Ft result in their axial polarization across cell interfaces and differential adhesiveness. We suggest that the role of Ds/Ft/Fj in long-range axially oriented planar cell alignment is a general function and that the regulation of the expression of its components would be crucial in the achievement of tissue uniformity in many other morphogenetic models or during tissue repair.

## INTRODUCTION

The tendency of densely packed structural units to align to each other is as a rule energetically (entropically) favorable and hence widely observed in nature (Beloussov, 2015). This property is often observed in multicellular assemblies, where it has major consequences for tissue morphogenesis. The establishment and maintenance of uniform long-range alignment patterns in tissues, based upon short-range interactions, is an important feature of development and is critical for correct physiological function (Gao, 2012) (Carvajal-Gonzalez and Mlodzik, 2014) (Hale and Strutt, 2015). Functional tissues not only require each cell acquiring its proper shape, size and polarity but also a stereotyped spatial multicellular organization. The particular alignment patterns of cells observed in tissues and organs such as the corneal stroma, vascular smooth muscle cells (SMCs), skeletal muscles or tendons are crucial for organ function (Lynch et al., 2003) (Holzapfel et al., 2005) (Boote et al., 2011) (Bourget et al., 2013).

In developing epithelia, cells maintain apico-basal polarity while progressively arrange in the epithelial plane, in many cases aligning along specific positional cues oriented with respect to the body axis. The temporal and spatial dynamics underlying the topographical organization, alignment and orientation of cells within large epithelia and the mechanisms driving these behaviors remain poorly explored. Short-range interactions seem to depend on cell-cell contacts, but long-range interactions only emerge as a result of coordinated intrinsic and extrinsic actions, in many cases via the generation of tensile forces amongst cooperatively aligned cells (Sugimura and Ishihara, 2013).

To explore the achievement of tissue-scale organization in growing epithelia, we analyzed the spatial-temporal progression of the long-range axially oriented alignment of histoblasts in *Drosophila.* The adult abdominal epidermis of *Drosophila* develops from four hemi-segmental iterated groups (nests) of imaginal diploid histoblasts embedded in between large polyploid Larval Epithelial Cells (LECs) (Madhavan and Schneiderman, 1977) (Ninov et al., 2007). Histoblasts are specified during embryogenesis and are quiescent all along larval development (Roseland and Schneiderman, 1979). At the onset of metamorphosis, and in response to a hormonal input, they start to divide and expand to replace the surrounding LECs, which become apoptotic (Madhavan and Madhavan, 1980) (Ninov et al., 2007). During their expansion, the histoblast nests preferentially stretch in the dorsal-ventral (D/V) direction through cell intercalation (Bischoff and Cseresnyes, 2009) (Umetsu et al., 2014). Cell divisions are not oriented along any preferential axis and apoptosis contributes little to the oriented nest expansion (Ninov et al., 2007) (Bischoff and Cseresnyes, 2009). Later, histoblast nests fuse to give rise to the anlagen of the adult epidermis (Garcia-Bellido and Merriam, 1971) (Roseland and Schneiderman, 1979).

At the end of metamorphosis, the abdominal epidermis of an adult fly consists of a monolayer of patterned cells polarized in the plane and an overlying cuticle (Casal et al., 2002) (Lawrence et al., 2002). By 50 h APF, most of the cells of the pupal epithelia have acquired an elongated hexagonal form and arranged in a uniform array of aligned rows oriented perpendicular to the A/P axis (see Fig. 1). How this precise long-range orientation is achieved is unknown and the cellular mechanism(s) and the genetic regulation underlying this behavior remain elusive.

**Figure 1.**
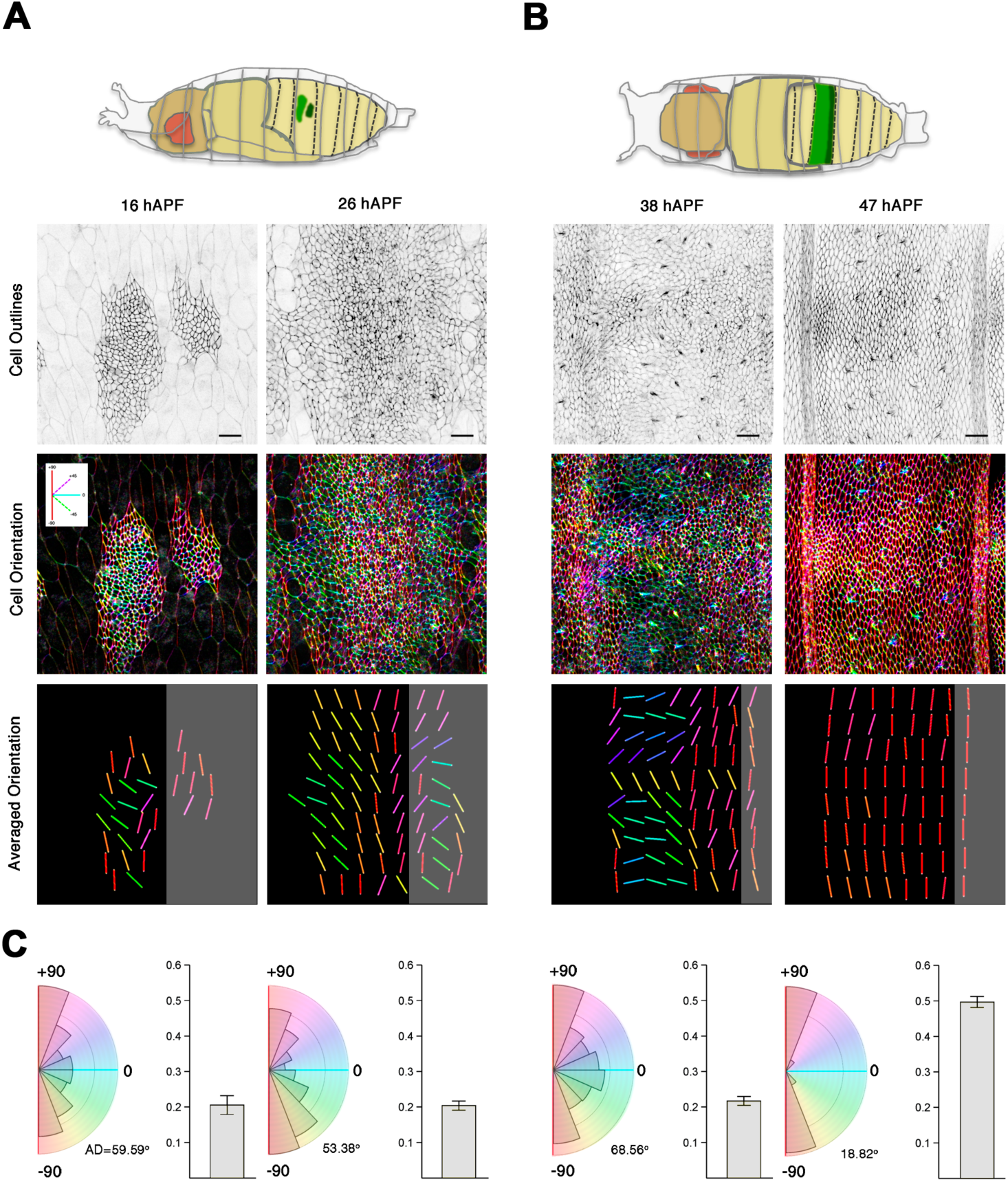
Axially Oriented Long-Range Alignment of Histoblasts. **A)** Histoblasts Expansion Stage. From top to bottom: Diagram representing in false colors the territory under study occupied by the dorsal anterior (light green) and posterior (dark green) histoblast nests of the third abdominal segment in the pupal abdomen at 16 h APF (dorso-lateral view); Grey inverted still images from snapshots of Movie S1 showing in-plane cell junctions (ZCL2207) at 16 and 26 h APF (beginning and end of the expansion period); Color-coded still images displaying the angular orientations of cell junctions referred to the A/P axis (inset displays the color code employed set in a way that red indicates alignment parallel to the A/P boundary, while cyan depicts perpendicular alignment); Color-coded orientation maps displaying the averaged short-range angular orientation of cell junctions for adjacent groups of neighboring histoblasts respect to the A/P axis (the A/P boundary corresponds to the apposition of anterior and posterior territories, shaded in black and grey respectively). **B**) Histoblasts Remodeling Stage. Equivalent images to A from top to bottom, except that the pupae diagram at the top it displays a dorsal view (38 h APF). Still images below correspond to the beginning (38 h APF) and end (47 h APF) of the remodeling period. See Fig. S1 and Experimental Procedures for detailed descriptions of the angular orientations color-code and ROIs analyses in A and B. Scale Bar is 22 μm. Anterior is to the left. C) Quantification of cell orientations relative to the A/P axis (left-rose diagrams) and cell alignment coherency (right-histograms) at 16, 26, 38 and 47 h APF (averaged for n = 6 pupae). The rose diagrams show the angular distribution of cell orientations with a binning size of 16° (different angles are color-coded as above). At each time point, wedge areas are proportional to abundance and the angular dispersion of the distribution is indicated by the angular deviation (AD) in degrees. Bar charts reporting the mean cell-cell alignment coherency at each time are bounded between 0 and 1. Mean values below 0.3 indicate averaged misalignment. Standard error of the mean (SEM) are displayed.

Directional morphogenesis is intimately linked to Planar Cell Polarity (PCP) (Gao, 2012), which is itself related to the activity of the Ds/Ft/Fj and the Core PCP pathways (reviewed by (Thomas and Strutt, 2012) (Sharma and McNeill, 2013a) (Matis and Axelrod, 2013) (Carvajal-Gonzalez and Mlodzik, 2014)). Of these pathways, the Ds/Ft/Fj has been shown to provide spatial cues orienting cell behaviors to tissue axes (Aigouy et al., 2010) (Bosveld et al., 2012) (Sharma and McNeill, 2013b). Ds and Ft are atypical cadherins that bind to each other heterophilically at cell interfaces (Strutt and Strutt, 2002) (Ma et al., 2003) (Matakatsu and Blair, 2004) (Brittle et al., 2012), while Fj is a Golgi kinase that phosphorylates both Ft and Ds, modifying their binding affinities (Clark et al., 1995) (Ishikawa et al., 2008) (Brittle et al., 2010) (Simon et al., 2010). It has been proposed that the positional output of the Ds/Ft/Fj pathway relies on the differential expression of its components (Simon, 2004) (Strutt, 2009) (Ambegaonkar et al., 2012) (Brittle et al., 2012) (Bosveld et al., 2012) (Hale et al., 2015) and *ds* and *fj,* indeed, are expressed in tissue-wide opposing gradients (Zeidler et al., 1999) (Zeidler et al., 2000) (Casal et al., 2002) (Yang et al., 2002) (Ma et al., 2003) (Simon, 2004) (Bosveld et al., 2012) while *ft* is uniformly expressed (Garoia et al., 2000) (Strutt and Strutt, 2002) (Ma et al., 2003) in various *Drosophila* epithelia. The Ds/Ft/Fj pathway regulates the asymmetric subcellular localization of the atypical myosin Dachs (Mao et al., 2006) (Brittle et al., 2012) (Ambegaonkar et al., 2012) amongst other targets.

Here, we examine the morphogenesis of *Drosophila* adult abdominal epidermis and show that the axially oriented long-range (uniform) planar alignment of the epithelia is an emerging phenomenon dynamically achieved in a long time-scale. Further, we found that progression through this process and the acquisition of an oriented pattern of cellular arrangements and cuticle structures is dependent on the function of the Ds/Ft/Fj pathway. We established that this pathway provides the positional cues necessary to bring the individual adjacent cells in register in a uniform orientation.

## RESULTS

### Histoblasts long-range directional order

To define how oriented cell alignment arises and evolves over developmental time during the morphogenesis of the *Drosophila* adult abdominal epithelial monolayer, we first performed a descriptive analysis using live imaging. We focused in the dorsal part of the abdomen where the adult epithelia derives from contralateral pairs of physically separated dorso-laterally located nests, one anterior (A) and one posterior (P), specified in different compartments of each segment (Kornberg, 1981) (Fig. 1A and 1B).

To monitor cell and tissue behaviors, we distinguished two sequential stages: First, the spreading of histoblasts at the expense of LECs (from 16 h to 26 h APF) until they envelop full hemisegments (tissue expansion) (Fig. 1A); Secondly, the planar reorganization of histoblasts (tissue remodeling) from full confluence (38 h APF) to the achievement of their final morphology (around 47 h APF) (Fig. 1B). Throughout both, expansion and remodeling, histoblast nests remain as a cohesive unit, displaying no gaps in the plane of the epithelium. We analyzed the dynamics of cell orientations both qualitatively and quantitatively by employing a GFP-tagged marker of cell junctions (ZCL2207) (see Movies S1 and S2) and the segmental A/P boundary as an axial reference (Fig. 1A and 1B). Locally averaged maps were extracted to highlight the progression in the orientation of cell alignments from a local to a global organization (see Fig. S1 and Experimental Procedures).

At the onset of the expansion stage (16 h APF), the histoblasts at the edges of the nests aligned mostly parallel to the A/P boundary, while those at the nests centre showed variable orientations (Fig. 1A). As expansion progressed, histoblasts arranged in groups oriented at different angles displaying local short-range alignment (Averaged Orientations, Fig. 1A - bottom panels). Cells located close to the A/P boundary showed A/P orientation, while at segmental contacts and within the nests they oriented mostly perpendicular to this axis.

At the initiation of the remodeling stage many histoblasts are already oriented parallel to the A/P boundary (Fig. 1B). Exceptional were the cells at the central region of the A nest, which were oriented almost perpendicular to the A/P boundary. As long-range remodeling proceeded, all cells realigned, sequentially from posterior to anterior, creating a uniform landscape (see Fig. S2). At about 47 h APF, all histoblasts had become uniformly oriented and aligned parallel to the A/P axis (Movie S1 and S2 and Fig. 1B, bottom right panel).

Quantitative analyses further substantiate that the long-range (uniform) oriented alignment of histoblasts was mainly achieved during the late remodeling stage (38 h–47 h APF, Fig. 1C and S2). Thus, the tissue coherence index increased from 0.2 to 0.5, *i.e.* the degree to which cells align to each other in a given direction (Fig. 1C).

Altogether, these analyses indicated that uniform axial orientation was an emerging phenomenon dynamically achieved over a long time-scale. Our results revealed that the long-range orientation progression followed a stereotyped A/P biased spatial-temporal pattern.

### The Dachsous/Fat/Four-jointed pathway is involved in the establishment of axial long-range cell orientation

Besides the obvious local influences that tissue boundaries and barriers may exert in orienting nearby cells (Nubler-Jung, 1987) (Umetsu et al., 2014) (Fagotto, 2015), in a wide sense, the cues that epithelial cells interpret and the mechanisms cells employ to implement global orientation are unknown. Considering the well-known role of the Ds/Ft/Fj pathway in defining the orientation of planar polarized cells with respect to tissue axes, we investigated its potential role setting the cues that could grant directional orientation guidance during abdominal epithelia morphogenesis. To explore this possibility we performed a spatiotemporal analysis upon perturbing the pathway activity.

In *ds (ds^UA071^/ds^38K^)* pupae, the spatial organization of the tissue was more irregular than in the wild type (*wt*). We found that by 26 h APF, at the end of the expansion phase, the anterior-ward progression of histoblasts re-orientation observed in the *wt* had not been taken place and just those cells adjacent to the A/P boundary were aligned to it (Fig. 2A, compare to Fig. 1A). By the end of the remodeling phase, at 47 h APF, most cells of the A compartment were organized in a patchwork of parallel-aligned cells deviating from the A/P boundary (Fig. 2A, see Movie S3). Only reached A/P orientation the cells located at the anterior border of the A compartment, at the A/P boundary or within the narrow P compartment (compare to Fig. 1B). In summary, cell alignment seemed to proceed properly at the local level but, overall, uniformity in cell alignment orientation (long-range organization) was not achieved (Fig. 2D and Movie S4).

**Figure 2.**
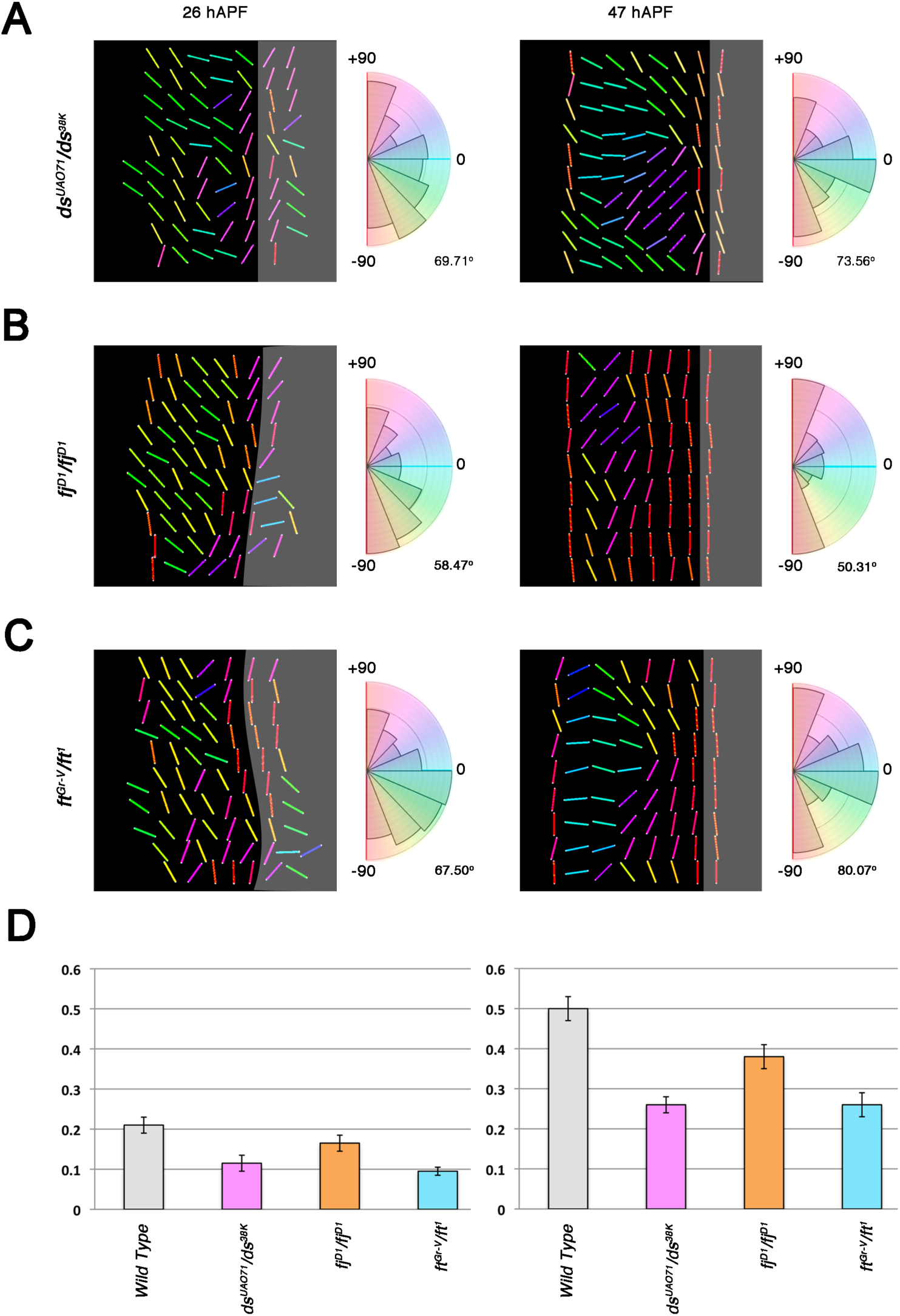
Alteration of the axially oriented long-range alignment of histoblasts in Ds/Ft/Fj pathway mutants. **A**) Color-coded averaged orientation maps and rose diagrams showing the angular distribution of cell orientations at the end of the expansion (26 h APF - left) and remodeling (47 h APF - right) stages for *ds (ds^UAO71^/ds^38k^)* mutant pupae (n = 6). B) Equivalent quantifications to A for *fj (fj^D1^/fj^D1^)* mutant pupae (n = 6). C) Equivalent quantifications to A and B for *ft (ft^Gr-V^/ft^1^)* mutant pupae (n = 6). As in Fig. 1, for the maps, the posterior area is shaded grey and the different angles for the rose diagrams are color-coded. Areas are proportional to abundance and the angular dispersion (AD) is indicated. Levels of significance were calculated with the Mardia-Watson-Wheeler W test. D) Averaged cell alignment coherency for *wt* (grey), *ds* (magenta), *fj* (orange) and *ft* (cyan) mutant pupae at 26 (left) and 47 h APF (right). As in Fig. 1, histograms report the mean cell-cell alignment coherency with values bounded between 0 and 1. Mean values below 0.3 indicate averaged misalignment. SEMs are displayed. Levels of significance were calculated with the Kolmogorov-Smirnov K-S test. Anterior is to the left.

Uniform orientation was similarly compromised in *ft (ft^Gr-V^/ft^1^)* and, to a lesser extent, in *fj* (*fj^D1^*) pupae (Fig. 2B and 2C). Patches of aligned cells with diverse orientations were observed at early stages in *ft.* These patches never resolved in a fully oriented pattern (Fig. 2D). In *fj* mutants, the initial cell orientation pattern was not markedly different from that observed in *wt* but full long-range orientation was as well affected and, in the most anterior part of the segment, uniformity was never reached. Overall, the *fj* phenotype was weaker than those of *ds* and *ft*.

Importantly, regardless of the consistent disruption of orientation uniformity in *ds, ft* and *fj,* we never observed defects in patterning or differentiation. In all conditions, the bristles-bearing cells (macro- and microchaete) arose in fairly normal positions *(e.g.* compare Fig. 3A and 3C) and their number was not altered suggesting that the Ds/Ft/Fj pathway had no major role in tissue patterning.

**Figure 3.**
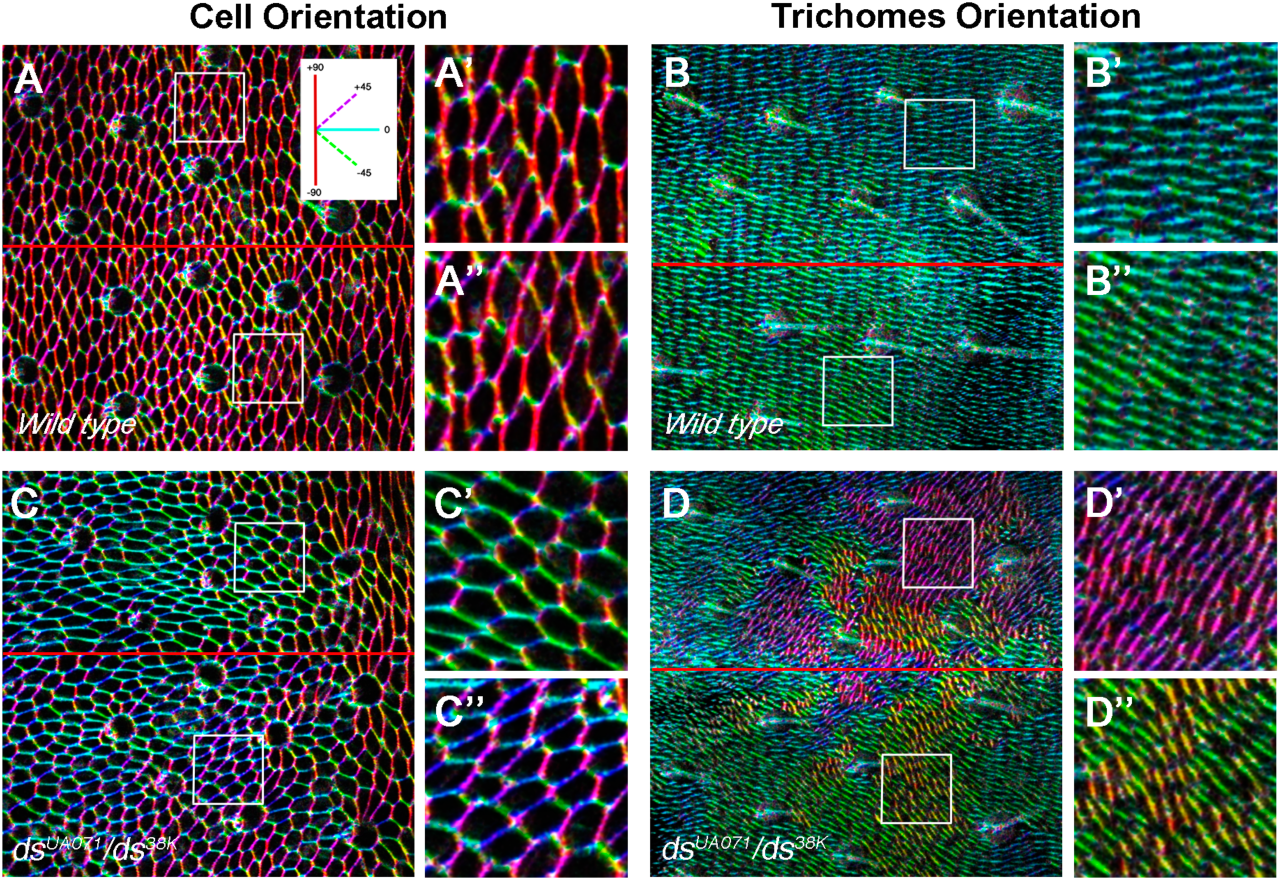
Long-range orientation and planar cell polarization are self-directed events. **A** and **B**) Color-coded angular orientations relative to the A/P axis for *wt* pupae (dorsal view) at 50 h APF of cell junctions (ZCL2207) **(A)** and actin (moesin::GFP) to visualize trichomes (**B**). Abdominal cells junctions display uniform orientation and align in rows along the A/P axis. Note that the *wt* trichomes uniform orientation is orthogonal to cells longer axis. Inset displays the color code employed: red indicates orientation parallel to the A/P boundary, while cyan depicts perpendicular orientation. **C** and **D**) Equivalent images to A and B but for *ds^UAO71^/ds^38k^* pupae. In this mutant condition, cells only reach short-range alignment and never get uniformly oriented along the A/P axis (see also Fig. 2). However, the orthogonal orientation of trichomes with respect to cells orientation is largely sustained in this condition. Primed panels (“and”) correspond to high magnification views of the boxed regions showed in the main panels. Scale bar is 12 μm. Anterior is to the left.

In summary, our data showed that the progression of the long-range orientation of cell alignments was severely impaired when the function of the Ds/Ft/Fj pathway was compromised.

### Long-range orientation and planar cell polarization are independent events

In the adult fly, epithelial cell planar polarity is manifested in the biased orientation of the trichomes (hairs) that decorate their apical surface, which, in the abdomen, invariably point posterior (Lawrence and Casal, 2013). For reasons that remain elusive (Zhu, 2009), *ds* or *ft* mutants display planar polarity phenotypes characterized by extensive trichome swirls in most of the A compartment (Casal et al., 2002) (Casal et al., 2006) (Matakatsu and Blair, 2006).

To explore whether the perturbed orientation of planar cell alignment found in *ds/ft/fj* pupae was related to the trichome swirling, we monitored hair formation in the abdomen of *wt* and *ds* pupae (Fig. 3). In the *wt,* around 44 h APF, once long-range orientation completion was almost achieved, multiple actin protrusions (up to 5) appeared at the posterior edges of each abdominal cell. These protrusions grew posterior and orthogonal to the A/P boundary (Fig. S3 and Movie S5) and their elongation was completed in about 6 hours. Importantly, we found that in *ds* mutants, although cells were no longer uniformly oriented (compare Fig. 3A to 3C), histoblasts were still largely polarized, trichomes arose at asymmetric positions within the cells as in the *wt* (compare Fig. 3B to 3D) and their outgrowth proceeded at normal speed (Fig. S3 and Movie S6). However, as the trichomes keep growing orthogonal to the long axis of each cell, abnormal cell orientations resulted in abnormal trichome patterns ((Fig. 3C and 3D, Fig. S3 and Movie S6). Strikingly, trichomes swirling patterns were not random but locally homogeneous (Fig. 3 and (Zhu, 2009)) drawing direct parallels with the stereotyped deviations from the A/P alignment observed for histoblasts orientations (see also Fig. 2).

Summarizing, the long-range orientation of the aligned epithelial cells preceded and was largely uncoupled from planar cell polarization. Trichome polarization within cells largely correlated with the organization of the underlying cell architecture, while their supracellular organization reflected the long-range orientation of the tissue.

### The expression of Ds, Ft and Fj is dynamically regulated during the abdominal epithelial morphogenesis

Our data suggested that, during morphogenesis, a correct input of the Ds/Ft/Fj signaling is essential for the directional long-range orientation of epithelial cells. It has been proposed that the positional output of the Ds/Ft/Fj pathway relies on the differential gene expression of its components (Simon, 2004) (Strutt, 2009) (Ambegaonkar et al., 2012) (Brittle et al., 2012) (Bosveld et al., 2012) (Hale et al., 2015). However, the spatiotemporal regulation of Ds, Ft and Fj expression during histoblasts morphogenesis remains unexplored and only the late patterns on pharate adults of *ds* and *fj* have been previously reported (Zeidler et al., 2000) (Casal et al., 2002). We therefore performed detailed expression analyses over time and space of Ds∷EGFP and Ft∷EGFP (protein traps) and fj-EGFP (enhancer trap) in pupae *in vivo*.

We found that from the initiation of histoblasts expansion, Ds was expressed in a tissue-wide axially graded fashion reversed in the A and the P compartments with maximum values at the A/P boundary. This pattern was sustained all throughout the early expansion and the late remodeling stages although the gradient steepness was progressively moderated (Fig. 4A and 4D and Movie S7). *fj* was expressed in a graded fashion opposing that of Ds from early stages. Its expression decreased from maxima at segmental borders towards minima at the A/P boundary (Fig. 4B and 4D and Movie S8). At early stages, *fj* expression was also higher dorsally, reaching maximum values at the presumptive dorsal midline. This D/V graded expression, weakened progressively overtime and eventually disappeared before the attainment of uniform axial cell orientation (compare Fig. 4B sequential panels). At that time, Ds and *fj* gradients were truly complementary and resembled those described for pharates (Zeidler et al., 2000) (Casal et al., 2002).

**Figure 4.**
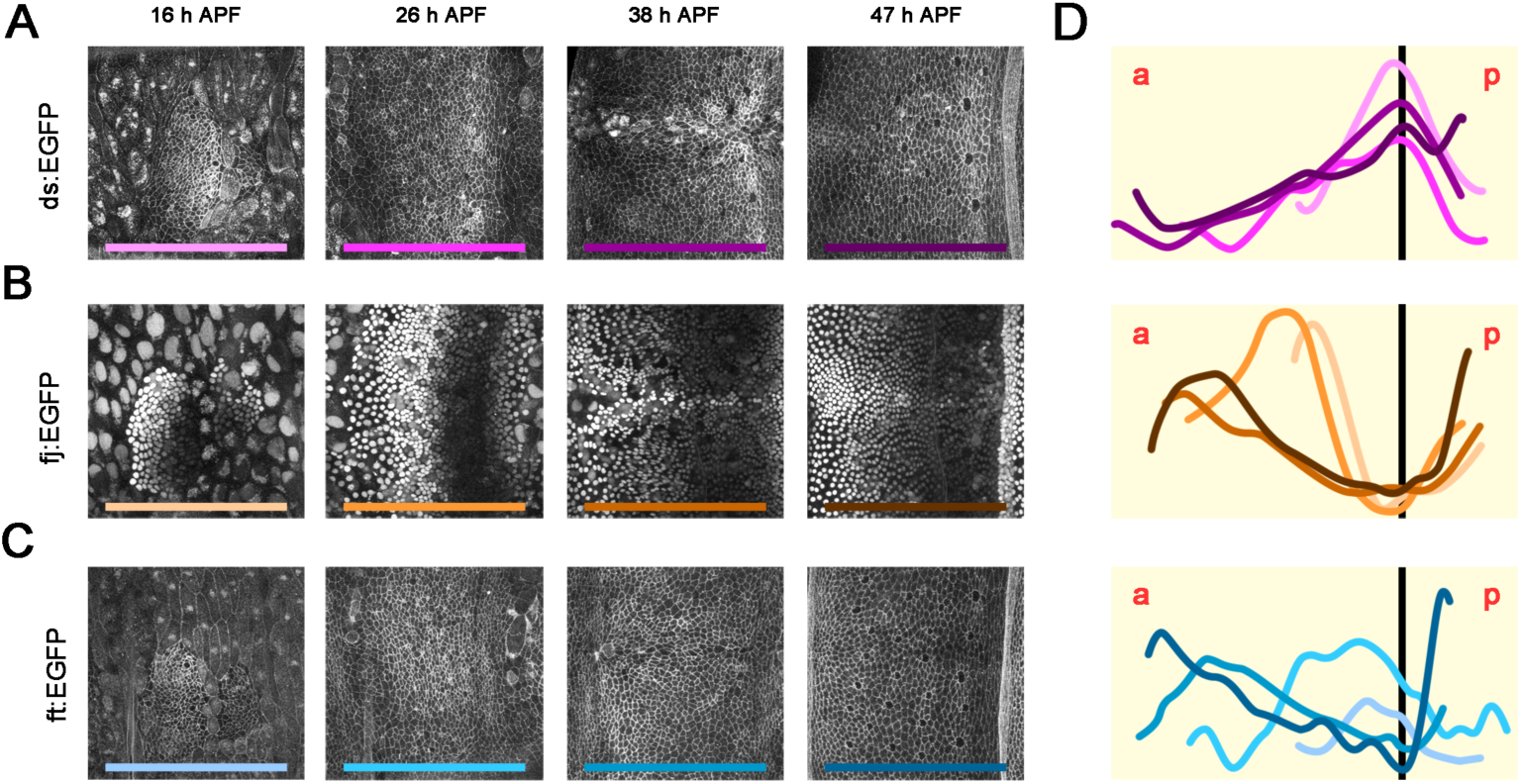
Dachsous, Four-jointed and Fat expression patterns. **A**) Expression pattern of Ds, visualized with a Ds::EGFP endogenous reporter during expansion (16 to 26 h APF) and remodeling (38 to 47 h APF). **B**) Expression pattern of *fj,* visualized with a *fj*-EGFP enhancer trap reporter (nuclear localization), at equivalent times as in A. **C**) Expression pattern of Ft, visualized with a Ft::EGFP. **D**) Graphs representing averaged expression profiles along the A/P axis in arbitrary units (see Experimental Procedures) for Ds (magenta-top), *fj* (orange-middle) and Ft (cyan-bottom) at the different time points displayed in A to C. Each profile kept its own scale along the A/P axis and they were aligned at the A/P compartment boundary (black vertical line). Profiles darkness for each marker was set to progress with aging from 16 (lightest) to 47 h APF (darkest). The color code is displayed under the original images in A to C. Scale Bar is 22 μm. Anterior is to the left.

Initially, in contrast to Ds and *fj*, Ft was not expressed in a graded manner. At the onset of expansion, Ft was spatially heterogeneous showing higher levels midway towards the posterior border of the A compartment decaying in both directions (Fig. 4D, lighter blue graphs). Ft expression progressively shifted anteriorly till, strikingly, delineated a gradient decreasing towards the posterior of the A compartment (Fig. 4C and 4D, darker blue graphs and Movie S9). Shortly before completion of axial long-range cell orientation, the Ft gradient became opposite to that of Ds, resembling *fj*.

### Long-range cell orientation responds to cues provided by the coordinated activity of the Ds/Ft/Fj pathway elements

Considering them together, the phenotypes observed in mutants for Ds/Ft/Fj pathway components and the spatiotemporal dynamics of their expression profiles, hint at an instructive role for this pathway in the establishment of long-range tissue orientation. To test if this function relied on a reciprocal regulatory activity between the pathway different components, we tested whether Ds or Fj were required for Ft expression remodeling and if this correlated with alignment orientation aberrations.

We found that in *ds* pupae, the final graded expression of Ft observed in the *wt* never developed (Fig. 5A, 5A′ and 5D). Ft levels remained highest at a middle position within the A compartment, decaying slowly in both anterior and posterior directions (Fig. 5B, 5B′ and 5D). This anomalous expression correlated with a failure on long-range tissue orientation, which was mostly affected on the anterior-central section of the A compartment (compare Fig. 5A″ and 5B″). Related defects in Ft expression and long-range cell orientation were also observed in *fj* pupae (Fig. 5C, 5C″, 5C″ and 5D). The impact of *fj* on Ft gradient formation and long-range orientation was, however, weaker than that of *ds*.

**Figure 5.**
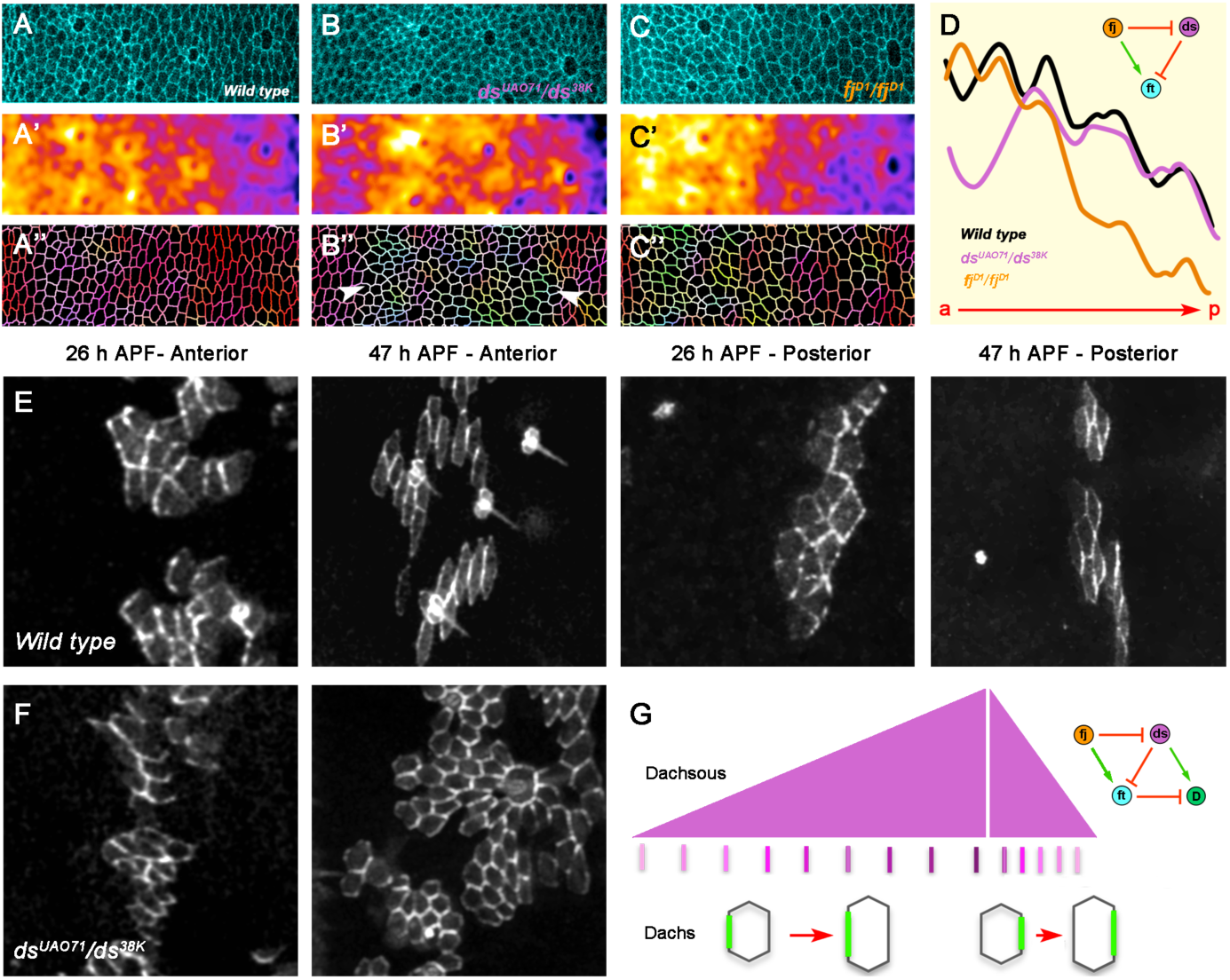
Regulatory crosstalk within the Ds/Ft/Fj network. **A-C**) Expression pattern of Ft (Ft∷GFP) along the anterior compartment of a third abdominal segment at 47 h APF in cyan false-color in *wt* (A), *ds (ds^UA071^/ds^38k^)* (B) and *fj (fj^D1^/fj^D1^)* pupae (C). **A′-C′**) Ft expression levels color-coded employing a "fire" LUT for the different genotypes (see Experimental Procedures). White-yellow indicates the highest intensity values while black-blue indicates the lowest. **A″-C″**) Color-coded segmented images from A to C displaying the angular orientations of cell junctions relative to the A/P axis. Multiple cells never properly align and misoriented ones topographically correlate with domains with inaccurate Ft expression (white arrowheads). Scale bar is 16 μm. Anterior is to the left. **D**) Graph representing the profile of Ft expression levels from the anterior to the posterior edge of the A compartment in *wt* (grey), *ds* (magenta) and *fj* (orange) backgrounds. The inset on the right highlights the functional relationships between the components of the Ds/Ft/Fj network. **E**) Clones of EGFP∷Dachs expressing cells in a *wt* pupae at 26 and 47 h APF (third abdominal segment). D is asymmetrically localized within cells at the membrane from early stages. At the A compartment D preferentially accumulates at the anterior side of each cell, while at the P compartment its location is reversed (white arrowheads). **F**) Clones of EGFP∷Dachs expressing cells in a *ds (ds^UA071^/ds^38k^)* pupae at 26 and 47 h APF (third abdominal segment). D expression is subcellularly asymmetrical but its preferential A/P axial localization is lost (magenta arrowheads). Some signs of axial orientation are still detectable even their overall directionality is randomized. Scale bar is 5 μm. Anterior is to the left. G) Diagram showing the asymmetric localization of D in *wt* cells (green) at the anterior and posterior compartments. D asymmetry relates to the A/P axis and lines up with the Ds gradient (magenta). The signaling network highlighted on the right represent the regulatory inputs of the Ft-Ds-Fj network onto D asymmetric localization.

Thus, a specific topographical and temporal balance amongst the different elements of the Ft-Ds-Fj pathway in terms of their expression/activity seemed to be essential for the completion of uniform alignment orientation.

How the cue(s) provided by the spatiotemporal distribution of the Ds/Ft/fj pathway activity could be executed was not clear. One of the best-recognized outcomes of the pathway is the subcellular polarization of the atypical myosin Dachs (D) (Mao et al., 2006) (Rogulja et al., 2008) (Ambegaonkar et al., 2012) (Brittle et al., 2012) (Bosveld et al., 2012). D polarization had been associated to uneven tension at cell cortices resulting into cell shape rearrangements (Bosveld et al., 2012). To explore whether the role of the Ds/Ft/Fj pathway on long-range orientation of cells alignment could be related to D polarization, we monitored small flip out clones expressing GFP-Dachs in transgenic flies in a *wt* background or in *ds.* In *wt* pupae, we found that D subcellular localization at cells edges was asymmetrically polarized from the onset of nests expansion, correlating with the *ds* A/P gradient at any position and time (Fig. 5E and 5G). D polarization at early stages was noisy, but became extremely robust as expansion proceeded. Remarkably, this uniform axis-guided polarized localization of D was affected in *ds* mutants (Fig. 5F), in which the Ds gradient was absent. D polarization linked up to the pseudo-randomized orientation of cell alignments that resulted from altering the combinatorial equilibrium between Ds, Ft and Fj.

Together, these results indicate that the abdominal epithelium became polarized very early and that D polarization was largely present at times and places in which the long-range orientation of the tissue had not been attained. D, and hence its contractile capabilities, did not seem to be causal for the long-range orientation of aligned histoblasts (see also Discussion).

### Ds/Ft/Fj directionally affects the local surface texture of the abdominal epithelia

To understand how the orientation of cell alignment is uniformly implemented and which is the nature of the directional cues mediated by the Ds/Ft/Fj signaling, we generated mitotic clones for *wt* and mutant cells for each component of the pathway, *ds^UAO71^, ft^G-rv^* and *fj^D1^* and compared, centering in the A compartment, their behavior *in vivo* at different scales, (see Experimental Procedures). At a large scale, we monitored different shape parameters: area, perimeter and aspect-ratio, which reflect the clones geometric proportions, and the main orientation angle that indicates their relation to the axial organization of the tissue. At a smaller scale we examined the clones radius of curvature (circularity and roundness). Last, at the smallest local scale, we evaluated clones roughness as an indicator of the ability of the cells of the clone to intermingle with their neighbors (solidity and convexity).

*wt* clones behaviors were stereotyped for all tested parameters and respond to the position of the clone within the tissue and to the position of the cells within the clones themselves (as we compared distinct parameters values between the anterior and the posterior halves/edges of the clones). We graphically described the collected parameters in color-coded cumulative plots (Fig. 6A and Fig. S4).

**Figure 6.**
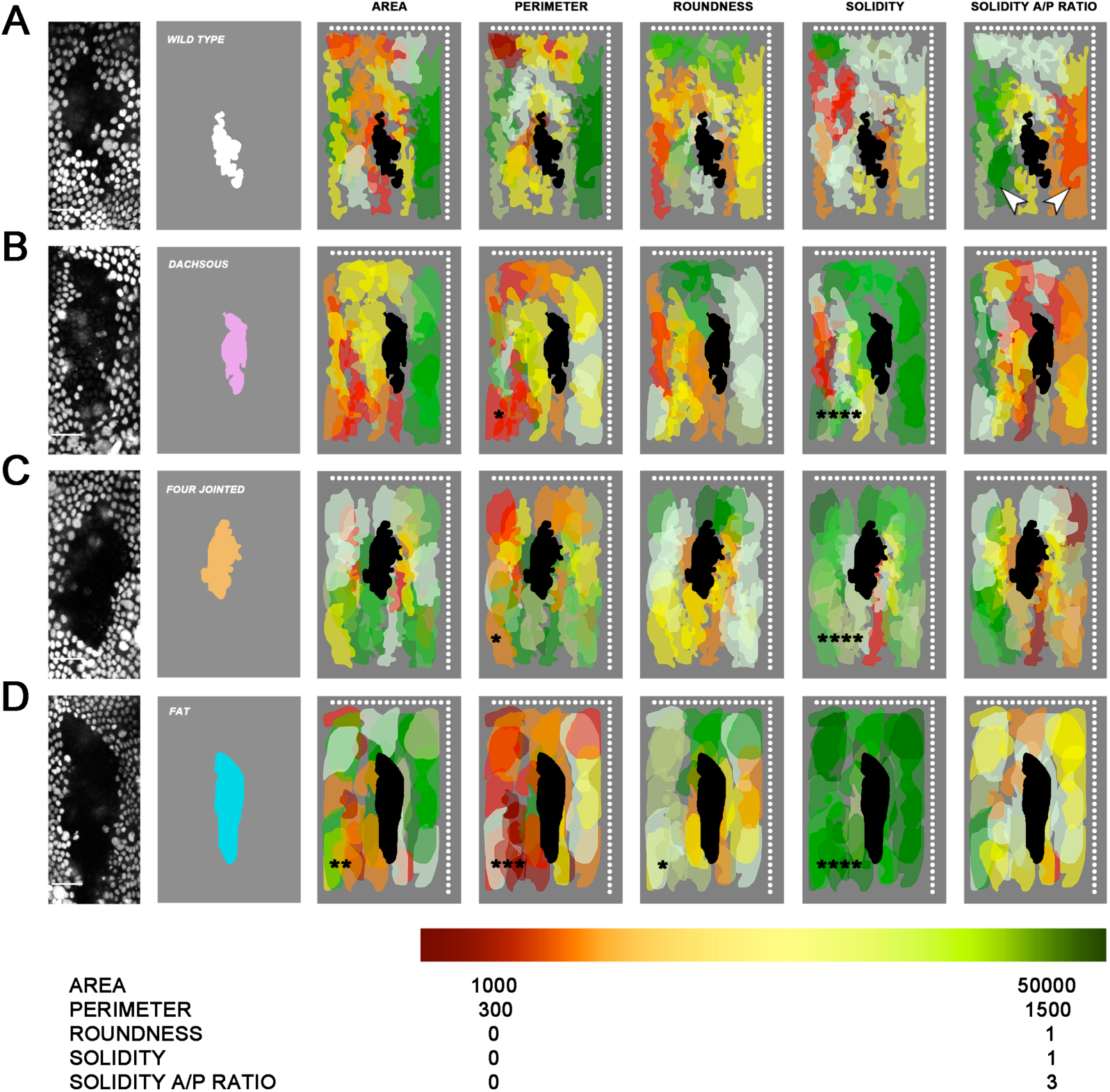
The Ds/Ft/Fj pathway affects tissue anisotropic expansion. **A**) *wt* clones. From left to right: Representative single clone of *wt* cells in a *wt* background (absence of RFP.nls expression); Mask corresponding to this representative clone (white) positioned within the anterior compartment; Cumulative topographic map displaying the shapes and positions of individual *wt* clones (n=26 - the representative clone is shown in dark grey) at the end of the remodeling phase (47 h APF) within the A compartment (dorsal and posterior edges are represented by dotted white lines). The values for geometrical (area and perimeter) and shape (roundness and solidity) parameters are represented by different colors according to a graded Green-Red LUT in sequential illustrations, from left to right. At the far right panel, the ratio of solidity between the anterior and posterior bisected halves of individual clones is also displayed. Arrowheads in this panel point to the different solidity axial bias between *wt* clones at anterior and posterior positions within the compartment. The different maximum and minimum values for each analyzed parameter in the graded Green-Red LUTs are stated at the bottom legend (see Experimental Procedures). **B**) *ds^UAO71^* clones (magenta). Equivalent panels as in A (n=26). **C)** *fj^D1^* clones (orange). Equivalent panels as in A (n=23). **D)** *ft^Gr-V^* clones (cyan). Equivalent panels as in A (n=27). Significant differences were evaluated by a Kolmogorov-Smirnov test. Black asterisks indicate significance levels for the averaged parameters values for each genotype vs. control *wt* clones (p*< 0.05, p** < 0.01, p*** < 0.001 or p**** < 0.0001).

*wt* clones showed low roundness, growing anisotropically in a dorso-ventral direction aligning with the A/P boundary (except those located close to the dorsal edge [see also (Garcia-Bellido and Merriam, 1971) and (Santamaria and Garcia-Bellido, 1972)]. They were also extremely rough from the beginning, with roughness increasing overtime. Interestingly, they showed a non-random directional bias progressively acquired over time in terms of solidity (Figs. 6A and Fig. S4). Most anterior clones showed posterior roughness and anterior smoothness at 47 h APF, while those located at posterior displayed the opposite.

In clones composed of mutant cells in an otherwise *wt* animal, we searched for topographically restricted defects in growth, anisotropy and orientation, and intercalation (cell intermixing and cell rearrangement) and their contribution to the final clone morphology (Fig. 6B to 6D). We found that the size (area) of mutant clones was just slightly larger than in controls and only significant in the absence of *ft* (in its anterior and central domain). The relative orientation of mutant clones was also unaffected by any genetic condition, all in all suggesting that the Ds/Ft/Fj pathway is not a primary participant in the control of growth or growth orientation. In contrast, clones of all genetic conditions, in particular of *ft*, showed significant perimeter reduction in comparison to *wt* ones (Fig. 6B to 6D). While the morphology (circularity or roundness) of mutant clones was not much different than that of *wt* ones (slightly more round for *ft*), the irregular borders observed in *wt* clones were lost and the complexity of the clones shape (solidity) was reduced for mutant conditions. The smoothness of the clones’ borders reflected decreased intermixing capabilities in between mutant and *wt* cells indicating that the major alterations in the morphology of mutant clones were the result of the tuning of cell-cell contacts at the clone borders. The reduction in clone shape complexity further indicated some sort of conflict between surface properties of mutant and surrounding *wt* cells. In this context, we found that the smoothness of borders was directionally biased depending on the mutated gene. While the surface of *ft* clones smoothed up all around their perimeter, *ds* clones smoothness was more prominent at their posterior edges, while for clones lacking *fj* this behavior was reversed. In close correlation with the distinct topographical influence of *ds* and *fj* in the long-range orientation of the tissue, *ds* clones asymmetric smoothness was observed anywhere within the compartment, while *fj* clones smoothness was restricted to anterior-central regions (compare Fig. 6B, 6C and 6D to Fig. 6A; see also Fig. 2).

Taken together, the data obtained from the clonal analyses showed that the smoothness of clonal boundaries and, hence, the suppression of neighbors exchanges of mutant cells, by distinct components of the Ft-Ds-Fj pathway is directional. This directionality was related to the expression pattern of the different components of the pathway and could be an outcome of the differential surface properties of mutant cells. Mutant histoblasts in clone borders will be unable to achieve correct heterophilic cell-cell interactions. The confrontation of different adhesive properties may generate surface tension inhomogeneities and these could, ultimately, trigger a failure in alignment orientation.

### Differential adhesion mediated by Ds/Ft/Fj leads to cell shape adjustments and rearrangements

To investigate the nature of the cellular activities implicated in the long-range orientation of cell alignment we determined the geometric parameters of individual cells at the time at which the tissue reached equilibrium. Cell geometries in an epithelial landscape can be accurately described by their interface contact angle and length (for cell edges shared between cells in contact), which are sensitive to the local balance between contractility and adhesion (Kafer et al., 2007) (Lecuit and Lenne, 2007). In *wt* pupae at 47 h APF, most histoblasts were elongated parallel to the A/P boundary and uniformly oriented (see also Figs. 1 and 3). Histoblasts contacted their neighbors at A/P oriented interfaces of 12 μm in length and at an open angle of 144° on average (See Fig 7A and 7E). Thus, as this anisotropical shape suggested, the surface tension and, most probably, adhesiveness were not uniformly distributed around the histoblasts perimeter at steady-state.

**Figure 7.**
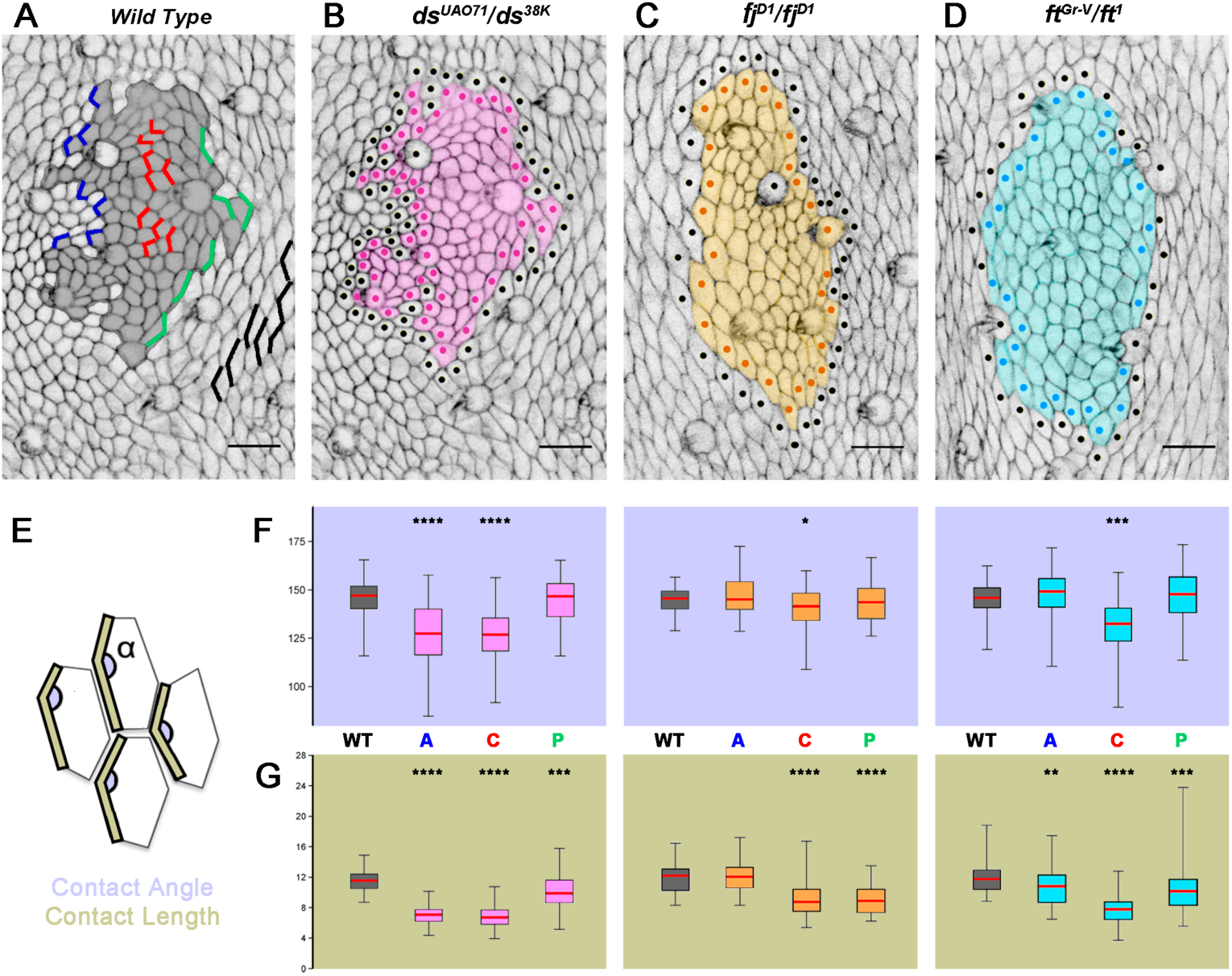
The Ds/Ft/Fj pathway modulates surface tension and long-range orientation of cell alignment. **A-D**) Mitotic clones induced by FLP/FRT-mediated recombination for *wt* (**A** - grey) *ds^UAO71^* (**B** - magenta), *fj^D1^* (**C** - orange) and *ft^Gr-v^* (**D** - cyan) in an otherwise *wt* background located at equivalent central positions in the A compartment of the third segment at 47 h APF. Cell outlines were labeled with ZCL2207. In **A** are represented the different cell contact edges described in E. WT - black lay outside the clone, A - blue are those at the clone anterior border, C - red are those within the clone and P - green those at the clone posterior border. For **B** to **D**, dots mark the cells at the internal and external clone edges. Scale bar is 14 μm. **E**) Diagram describing the parameters employed to estimate the relative mechanical (adhesion/contractile) directional properties from cell shape geometries. The contact length (light brown) corresponds to the length of the perimeter shared between cells along their A/P oriented contacts. The contact angle (α) (light purple) reflects the degree at which they contact. **F-G)** Range of contact angles **(F)** and contact lengths **(G)** for each genetic condition *ds^UAO71^* (magenta), *fj^D1^* (orange) and *ft^Gr-V^* (cyan) represented with box and whiskers plots. The box size indicates the range of contact length and angles in the central 50% of the distribution (between the 75 and 25 % quartiles). The red lines within the boxes indicate the median values. Whiskers extend to the minimum and maximum values. Two-sample unpaired two-tailed student t-test for significance was applied and distributions significantly different from WT were marked (p*< 0.05, p** < 0.01, p*** < 0.001 or p**** < 0.0001). WT =75, C = 132, A = 69 and P = 53 contacts in 5 *ds* clones; WT = 108, C = 191, A = 62 and P = 74 contacts in 7 *ft* clones; and WT = 45, C = 94, A = 40 and P = 47 in 4 *fj* clones).

We then measured cell shape geometries within, outside and at the anterior and posterior edges of clones lacking *ds, fj* or *ft* (Fig. 7B to 7D). Histoblasts located at a distance of mutant clones all have contact angles and lengths values equivalent to those of *wt* animals (Fig. 7E to 7G). However, for any mutant condition, those histoblasts located at the centre of clones displayed significantly smaller contact angles (approaching 120°) and interfaces lengths (particularly for *ds* and *ft)* than those outside the clones. This indicated that in the absence of components of the Ds/Ft/Fj pathway cells geometrical anisotropy and adhesiveness were autonomously affected.

Notably, differences were also observed for clones’ anterior and posterior boundary cells (those cells in the clone in contact with the surrounding *wt* cells) and their analysis allowed to assess directional effects on cells geometry for the different pathway components. They also let study their correlation to clones edges smoothening. For *ds* and *fj* mutant clones, the contact angles between background *wt* and clones edge cells were different anteriorly and posteriorly (Fig. 7F). Averaged contact angles at the smooth sides of the clones (posterior for *ds* and anterior for *fj*) were alike to those between *wt* cells, while at their rough edges, contact angles were similar to those amongst mutant cells. For *ft* clones with isotropic smoothness, averaged contact angles at both sides, anterior and posterior, were identical to those between *wt* cells (Fig. 7F). The averaged lengths of the contacts at the interfaces followed the same trend as contact angles, except for *ft*, where they were, irrespectively of their position at the clone edges, of an intermediate length to those between mutant cells and those between background *wt* cells (Fig. 7G). Thus, it appeared that *wt* cells with an intact Ds/Ft/Fj signaling network could directionally rescue the shape, alignment and orientation of mutant cells at short distance.

## DISCUSSION

Investigating how cells topographically organize and segregate in different morphogenetic processes has yielded many insights into the diversity of cellular behaviors that contribute to the shaping of tissues and organs. Regarding to the morphogenesis of planar epithelial cell/tissues, studies in multiple model organisms have revealed significant commonalities and point to the Planar Cell Polarity (PCP) pathways as key coordinators and to mechanical forces as essential partners (reviewed in (Heisenberg and Bellaiche, 2013) (Devenport, 2014). By applying *in vivo* analyses to the genetically and morphologically tractable morphogenesis of the adult abdominal epidermis of *Drosophila* (Ninov and Martin-Blanco, 2007), we have characterized the stepwise long-range organization of the tissue and uncovered a new mechanism for its control. Prevailing views emphasize the activities of motor proteins and cortex contractility in directing epithelial cell and tissue rearrangements (Lecuit et al., 2011) (Heisenberg and Bellaiche, 2013). However, we found that shaping the epithelial landscape during histoblasts expansion mostly respond to an adhesion code mediated by the Ds/Ft/Fj pathway topographically balanced along the A/P axis (Fig. 8). The directional cues dictated by the pathway put the epithelial cells on the right track, orienting their changing shapes (see below).

**Figure 8.**
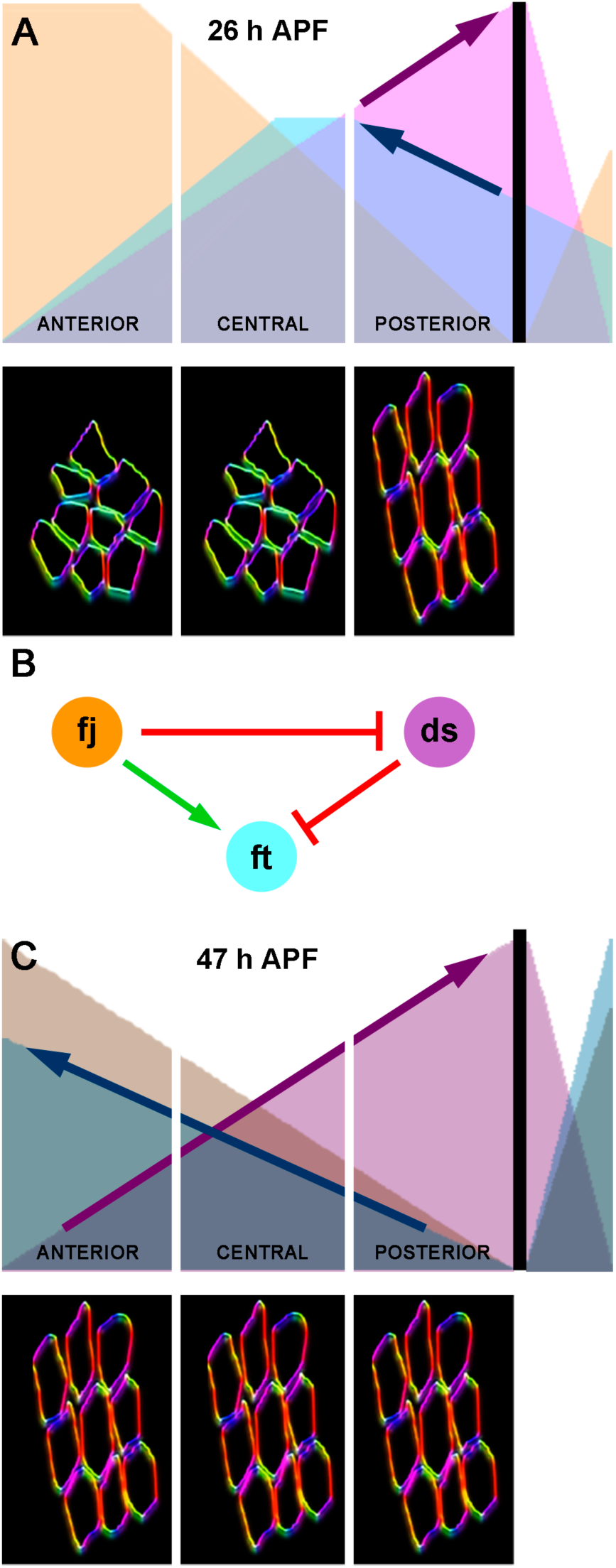
The Ds/Ft/Fj pathway coordinates long-range orientation of cell alignment along the A/P axis. **A**) At the onset of expansion, histoblasts display heterogeneous cell shapes and orientations. Progressively, they locally align to each other in small patches of cells with common orientation (end of expansion). At this time the expression patterns of Ds (magenta), *fj* (Orange) and Ft (Cyan) are uneven (top) and a reverse gradient of Ds and Ft is only found at the most posterior domain of the A compartment. This domain correlates with the most precocious area showing long-range cell alignment orientation (bottom). **B)** Reciprocal regulatory contributions between the different members of the Ds/Ft/Fj network, mainly the negative influence of Ds on Ft, results in the refinement of this landscape, and in particular on the establishment of a compartment-wide Ft gradient. **C)** In this way, the expression patterns of Ds, *fj* and Ft at the end of remodeling reach their final shapes and their activities get balanced along the A/P axis. The establishment of a compartment wide inverse relationship between Ds and Ft (top) implements over time a specific surface tension field through differential adhesion, favoring the stability of A/P oriented cell contacts. We propose that differential adhesiveness provides the necessary cues for the establishment of the long-range axially oriented planar alignment of histoblasts during abdominal morphogenesis (bottom). This proposition is supported by epistatic analyses, local traits of clonal behaviors and autonomous and non-autonomous alterations in cell-cell contacts geometries.

Shaping tissues may involve many different cellular and molecular activities. Coordinated cell-cell rearrangements trigger tissue re-orientation (Aigouy et al., 2010) (Suzanne et al., 2010) but can also lead to convergent extension (Keller, 2006). Alternatively, the apical constriction of epithelial cells may direct local bending of epithelial sheets (Pilot and Lecuit, 2005). Further, spatially controlled cell proliferation and growth, cell division orientation, and cell death can also give rise to global tissue changes (Hopyan et al., 2011) (Mao et al., 2013) (Campinho et al., 2013) (Legoff et al., 2013). We found that the axially oriented long-range alignment of histoblasts emerges in a precise spatio-temporal manner from initially naive growing epithelial cells with dispersed orientation (Fig. 1). We also found that their alignment proceeds through coordinated and oriented changes in cell shape and probably depends on the oriented activity of the Ds/Ft/Fj pathway along the A/P axis. This pathway implements over time a dynamic and stereotyped differential adhesion code axially oriented throughout the epithelium. When the expression of *ft, ds* or *fj* gets critically weak, the relative orientation of cell shapes is severely disturbed. This occurs without alterations in tissue patterning or cell differentiation.

It is known that differential adhesive properties between neighbors prevent cell intermingling, which in clones, to minimize the contacts between cells that are not alike, lead to borders smoothness (Nardi and Kafatos, 1976) (Nubler-Jung and Mardini, 1990) (Lawrence et al., 1999). Notably, we found that in the developing abdominal epithelia, major differences in roughness, perimeter and, to a lesser extent, on roundness are developed in mutant clones for members of the Ds/Ft/Fj pathway (Fig. 6). For each component, the edges of the clones get smoother than for *wt*, particularly for *ft* where clones were smooth all around their perimeter. Smoothness was also evident for *ds* and *fj*, although in these cases, it showed a directional bias. Smoother edges were found at the mutant clones’ borders facing the flank of the A/P axis showing maximum expression of the affected gene: anterior for *fj* and posterior for *ds.* Altogether, these data support a role for the combinatorial activity of the Ds/Ft/Fj pathway as the tissue proceeds through long-range orientation in generating directional information at cell junctions (Fig. 5). These directional inputs lead to planar conflicts where cells have different adhesive properties, and these would result in clones with smooth borders at specific edges, as we observed.

We also found that the early subcellular polarization of *D* in response to the Ds/Ft/Fj pathway and the long-range orientation of cell alignments are uncoupled. *D* polarized subcellular distribution was sustained in *ds* but re-oriented analogously to the changes of cell shapes observed. Further, *D* loss of function in clones in an otherwise *wt* background did not affect the axial orientation of cell alignments (Mao et al., 2006) (Mao et al., 2011) or cell shape (Legoff et al., 2013). Therefore, a bias in contractility at the cell cortex that could be generated by the asymmetric distribution of *D* seems dispensable for long-range tissue orientation. It is thus tempting to propose that the differential adhesion through heterophilic interactions between Ds and Ft may be at the root of the cues employed by histoblasts to axially orient in the long-range. This differential adhesion code would affect the capacity of the cells to change shape and exchange neighbors and to coordinately orient to maximize contacts (Fig. 8). Indeed, we found strong deviations at clones’ edges or within clones in mutant conditions from the oriented geometrical bias of *wt* cells at steady-state (Fig. 7). Contact angles and contact lengths between cells seem to specifically adjust depending on the clonal genotype and the relative location of the clone within the compartments.

An evolving pattern of differential adhesion at cell junctions would have a direct input into surface tension (Kafer et al., 2007) (Lecuit and Lenne, 2007) and therefore on cell and tissue shapes. Assuming that cells and tissues tend to minimize their surface free energy (Foty and Steinberg, 2005), the binding of adhesion molecules, causing cells to spread their cell-cell contacts, and the contractile activity of the actomyosin cell cortex counterbalancing adhesive forces, will be key determinants of cell and tissue shapes (Amack and Manning, 2012). Tensile patterns rather than cell positions would therefore play instructive roles in the acquisition of long-range morphological order. Importantly, they, at any given point in time, will reflect the recent developmental history of the tissue (Beloussov, 2015). Along this line, during the expansion and remodeling of the abdominal epithelia, the expression pattern of Ft modulated by Ds and *fj* evolves into an A/P gradient spanning whole compartments (Fig. 4). This expression refinement will result in the spreading, all over the epithelia, of a counterbalanced adhesion share between Ft and Ds that will delineate an axially oriented surface tension landscape instructive for long-range alignment. The most precocious areas to orient, which are located at the posterior edge of the A compartment would correspond to those domains where the compensate Ft A/P gradient originates (Figs. 1 and 4). Cell aligning orientation would then proceed anterior along the body axis following the spreading of the tissue-wide Ft gradient.

The role we uncovered of the Ds/Ft/Fj pathway implementing the uniform orientation of the alignment of histoblasts that, does not relate to any developmental input involved in patterning, cell specification and/or differentiation. The directional evolution of the expression of the different Ds/Ft/Fj pathway elements sets up a spatially and temporally controlled differential adhesion code between Ds and Ft. This provides the basis for the establishment of a dynamic surface tension “epipattern” unfolded over time that directs cell shape changes and cell intermingling, ultimately guiding the orientation of the long-range alignment of the epithelial cells.

## EXPERIMENTAL PROCEDURES

### Fly Stocks and Genetics

The mutant alleles employed *(ds^UAO71^, ds3^8k^, ft^G-rv^, ft^1^, fj^D1^)* are all thoroughly described in Flybase (flybase.org). Transgenic stocks carrying GFP trap constructs employed in this work from FlyTrap (flytrap.med.yale.edu) include w; P{w^+mC^=PTT-un1}*ZCL2207*, inserted in the ATPα gene that labels septate junctions as a cellular marker and *fj^CB04634^*, inserted in the regulatory region of *fj.* Other markers include Moesin∷GFP *(sqh* promoter driving the expression of a fragment of Moe fused to GFP), ubi-RFP.NLS FRT40A and FRT42D ubi-RFP.NLS, all available from the Bloomington Stock Center. The Ds∷EGFP and Ft∷EGFP endogenous reporters and the Act5c>stop>EGFP∷Dachs transgenic flies were kindly provided by David Strutt (Brittle et al., 2012).

*ds,* and *ft* viable mutant flies were generated by combining two different alleles of *ds (ds^UAO71^* with *ds^38K^)* and two different alleles of *ft (ft^G-rv^* with *ft^1^),* respectively. The null allele *fj^D1^* is homozygous viable. In order to visualize cell outlines, trichomes orientation or Ft expression, the different markers were incorporated in each mutant background.

The genotypes of the studied mutants were as follows:

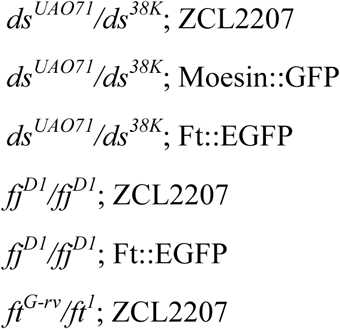

Flies husbandry was performed at standard conditions.

### Live Imaging of the Pupal Abdominal epithelium

Pupae were staged according to Baindbridge and Bownes (Bainbridge and Bownes, 1981) and timed employing the puparium formation as a reference (hours after puparium formation - h APF). Pupae were dissected and mounted as previously described (Ninov and Martin-Blanco, 2007), with the exception that the whole pupal case was removed before imaging. All images and time-lapses monitored, alternatively, the dorso-lateral expansion of ipsilateral A and P nests or the mid-dorsal fusion of contralateral nests. We preferentially focused on the third dorsal abdominal segment because its size, shape and differentiation pattern is almost identical in both females and males.

Pupae were imaged on an inverted Zeiss LSM 700 confocal microscope, with a Zeiss Plan-Neofluar 40 X/1.3 (NA) oil immersion objective lens at 0.7 X zoom. Between about 16 and 32 slices for each image were recorded, with a z-stack step size of 1 μm, resulting in a voxel size of 0.22 X 0.22 X 1 μm for a total of 1024 X 1024 pixels in each slice (resolution of 4.479 pixel per μm). This image acquisition set up was used for both, single image or time-lapse captures. The time interval for each time-lapse recording was set at 5 minutes. Such time resolution is suitable enough to extract relevant details of tissue expansion and remodeling with high fidelity.

Laser intensity was kept to a minimum and no averaging was taken to reduce acquisition bleaching and to minimize photo-toxicity. Pupae were kept in culture after imaging, and more than 95% hatched normally, suggesting negligible tissue damage during image acquisition.

### Image processing, analysis and statistics

ImageJ (imagej.nih.gov/ij/) was used for viewing and processing all confocal images and time-lapses. All images and movies are maximum projections of Z-stacks, unless otherwise stated. To minimize undesirable fluorescence noise from circulating macrophages, the number of Z slices employed into 2D tissue reconstruction was kept into a minimum.

To assess planar cell alignments and orientation we first set internal tissue landmarks to define a planar coordinate system. As main landmark we employed the A/P compartment boundary tilted to fit a vertical axis defining anterior (left) and posterior (right) domains. In this way, all the cell edges oriented in line with the A/P boundary were defined as aligned along the A/P axis of the segment. Conversely, each position perpendicular to the A/P boundary corresponded to the horizontal axial coordinates. All cell edges oriented perpendicular to the A/P boundary were defined as aligned along the D/V axis of the segment.

Tissue orientation was monitored with Orientation J (Fonck et al., 2009) (Rezakhaniha et al., 2012), an ImageJ plug-in based on structure tensors that maps the angle of oriented structures in input images. Considering that cell edges are indeed oriented structures, we used this plug-in to follow and measure the degree of planar cell alignment and its orientation. Briefly, the orientation of a cell can be described by the angles (axial range between −90 and 90 degrees) of their edges with respect to defined planar coordinates. To represent individual cell edge orientations with respect to our planar coordinate system, the graded color code generated by the plug-in was set in a way that red color indicated cell orientation parallel to the A/P boundary (−90° and 90°), while cyan depicted cell orientation perpendicular to the A/P boundary (0°).

To generate maps of local-averaged cell edge orientations, the histoblast territory was manually subdivided in small adjacent and not overlapping Regions of Interest (ROIs) of uniform weight (64 × 64 pixels, which approximately correspond to 20 × 20 μm). For each ROI, the resultant local predominant cell edge orientation corresponding to the direction of the largest eigenvector of the tensor was determined. These were plotted in locally averaged orientation maps centered at each ROI position as color-coded bars describing their mean angle.

The global alignment orientation of the tissue was calculated by extracting the local-averaged orientations of individual ROIs determined from six individual pupae for each condition and time and distributing their values in 16 sequential bins covering 360°. For simplicity, the distribution was represented in rose diagrams (180° spanning from +90° to −90° with respect to the A/P boundary) covering a hemicircle. For each class, abundance was represented as proportional to its area.

The strength of cell alignment along the calculated dominant orientation can be expressed by a coherency index defined as the ratio between the difference and the sum of the largest and smallest tensor Eigen values (Fonck et al., 2009; Rezakhaniha et al., 2012). Coherency is bounded between 0 and 1, with 1 indicating highly oriented structures and so uniform cell alignment with respect to the calculated dominant orientation and 0 indicating isotropic areas with low degree of alignment. Averaged values of coherency for six individual pupae for each condition and time were reported in bar charts.

Topographical information on axial alignment through the epithelial landscape (Fig. S2) was determined for the A compartment domain during the remodeling phase. On equivalent specific 2D ROIs from images selected at different time points we sequentially performed several tasks employing different Image J plug-ins: reslicing along the A/P axis; sum of slices projection; RGB color channels split; selection of a 1D ROI along the axis; and generation of plot profiles for the full and red color channel signals. The ratio between the red (indicative of cell alignment parallel to the A/P boundary (−90° and 90°) and the full signal plot at each point let to normalize the values of the average orientation level of cell edges at each position along the chosen axis (A/P) for the selected ROIs.

Expression profiles along the A/P axis of *ds, fj* and *ft* at each stage representing averaged pixel intensities across rectangular selections covering the territory occupied by the nests were plotted in 2D graphs using the Plot Profile Option of ImageJ.

All graphs and associated statistics were performed with PAST software (Hammer et al., 2001). The statistical analyses employed for each set of data are listed in each figure legend. Error bars represent standard error of the mean (± SEM) and statistical significance was determined using unpaired two-tailed Student’s t-test for equal mean or the non-parametric Kolmogorov-Smirnov K-S-test. For axial data we have performed circular statistics, calculating the mean Rayleigh (R) vector and the angular deviation (AD) for each distribution. The Mardia-Watson-Wheeler W-test was then used to compare the resultant distributions.

In all the cases the level of significance as been set as: p* ≤ 0.05; p** ≤ 0.01; p*** ≤ 0.001; p**** ≤ 0.0001; NS, not significant (p > 0.05 compared to control).

### Clones Generation

*ds^UAO71^*, *ft^G-rv^* and *fj^D1^* mutant clones, as well as *wt* control clones, were generated using the FLP/FRT mediated mitotic recombination technique (Golic and Lindquist, 1989) (Xu and Rubin, 1993) and marked by the absence of a nuclear-tagged Ubiquitin-Red Fluorescent Protein. ZCL2207 chromosome was incorporated in the appropriate stocks to mark cell outlines. Mutant clones were induced by heat-shock treatment of third instar larvae in a water-bath at 37°C for 1 hour. Upon clone induction, living pupae were dissected and mutant clones identified and imaged at desired stages. We analyzed clone parameters at nearly the end of expansion (26 h APF) and at the end of remodeling (47 h APF) in parallel experiments.

Genotypes for mutant clones in *wt* background with labeled apical cell outlines:

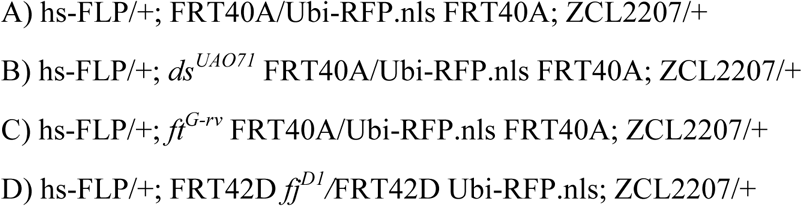

To create small clones of EGFP∷Dachs expressing cells in *wt* or *ds^UAO71^/ds^38K^* backgrounds, transgenic flies carrying the stop cassette Act5c>stop>EGFP∷Dachs were crossed with flies carrying a hs-FLP transgene and heat shocked for 6-8 minutes at 37°C at third instar larvae. Living pupae were dissected and the EGFP∷Dachs expressing cells identified and imaged at the desired stage.

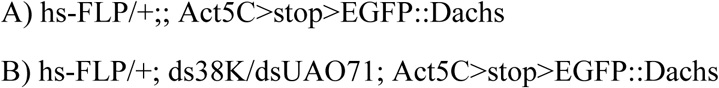

### Clonal Analysis

To infer geometrical and shape parameters from clones of cells, we first identified the patches of cells devoid of the RFP.nls marker from individual images collected at the same conditions. We manually segmented the border of individual clones and topographically defined the location of their centroids relative to specific reference points, the A/P boundary and the dorsal midline. All clones were cumulatively superimposed onto diagrams representing the third abdominal hemi-segment. The resultant cumulative clone diagrams for different stages and conditions were scaled to allow the comparison of eventual differences in clone orientation and shape.

From the segmented clone outlines we extracted geometrical and shape parameters using ImageJ.

**Area** = area of particle (in square pixels),

The clone area is the projected two-dimensional area calculated as the sum of the areas of each individual pixel circumscribed within the clone boundary

**Perimeter** = length of the clone perimeter in pixels.

The perimeter of the clone is defined as the total length of the clone boundary.

**Aspect-Ratio (AR)** = major axis/minor axis.

Aspect-ratio is defined as the ratio of the major axis to the minor axis of the best-fit Legendre ellipse inscribed into the clone area. Thus, as the length and width of the shape approach the same value, the aspect ratio approaches 1. The converse of this is also true. Thus, as the measurements of the length and width of the shape diverge, the aspect ratio approaches infinite.

**Angle** = Angle of the clone main axis with respect the A/P axis.

The angle between the major axis of the clone and the axial references was used to estimate the relative orientation of the clone with respect to the tissue boundaries

**Circularity** = 4π*area/perimeter^2.

Circularity is a shape parameter sensitive to both clone form and surface roughness, since it measures the deviation of a particle from a circle that can either result from either changing particle elongation or roughness. As a shape becomes more round and smooth, the circularity would approach 1. Conversely, as a shape becomes less round or as the shape becomes less smooth, the circularity should approach 0.

**Roundness** = 4*area/(π*major_axis^2)

Roundness is a shape parameter that measures of how closely the shape of a clone approaches that of a circle. It is just sensitive to clone form. Values oscillate between 0 and 1, with 1 indicating almost circular shapes

**Solidity** = area/convex area.

Solidity is the measurement of the overall concavity of the clone area. It is defined as the clone area divided by the convex hull area. The convex hull area/perimeter is defined as the smallest convex area/perimeters that contains the shape of the clone. Solidity describes area-based roughness. Values close to 1 indicate that the clone have a convex shape with smooth and straight borders. Increasing irregularities (indentations) in the clone outline decreases this value toward 0.

**Convexity** = Convex perimeter/Perimeter

Convexity is a measurement of the clone edge roughness. When a shape becomes rougher, the increment of its perimeter depends on the size and number of the irregularities in the shape. Thus, as the shape becomes less smooth, the convexity approaches to 0. Conversely, as the shape becomes smoother, the convexity approaches to 1.

The ratio for shape parameters (circularity, roundness, solidity and convexity) between the anterior and posterior edges of each clone was calculated by bisecting the clones’ masks along their A/P axis. Shape parameters values were calculated independently for each hemi-clone. A simple ratio between the two values was determined. Values approaching 1 indicated no differences between the anterior and posterior edges, while above 1 denote anterior higher values and below 1 posterior higher values.

For every parameter and ratios an independent red to green color-code for increasing values was manually created. This color-code was consistently applied to every clone and the coded values were applied to the cumulative clones images presented in Figs. 6 and S4.

### Contact angle and contact length measurements

To estimate the relative interfacial tension from cell shape geometries (Hayashi and Carthew, 2004) (Kafer et al., 2007) we incorporated the ZCL2207 marker into the background of the stocks employed to generate clones. Cell shape geometry at the steady-state (47 h APF) can be described by two parameters: the contact angle and the contact length between cell edges (cell interfaces) (Fagotto, 2014). The balance between contractile and adhesive strengths at the interface between two adjacent cells defines the cell interfacial tension (Hayashi and Carthew, 2004) (Kafer et al., 2007) (Rauzi et al., 2008). At the steady state, the abdominal cells have a polygonal shape approximating a hexagonal form and are packed in A/P oriented contiguous rows. The measured contact angles and lengths correspond to those cell edges facing adjacent rows aligned along the A/P axis. Thus, we have assessed the interfacial geometry between A/P oriented rows. The averaged contact angle and contact length were then calculated for cells within, at edges and outside the clone areas. Mutant cells in contact with *wt* cells at the anterior (A) or the posterior (P) side of each clone were analyzed separately.

## ACKNOWLEDGEMENTS

We thank Antonio García-Bellido for insightful discussions. We also thank the members of the EMB laboratory for continuous support and encouragement. We are grateful to Marco Milan, Florenci Serras, Nic Tapon, Francois Graner and David Strutt for reading earlier versions of this manuscript and to numerous members of the *Drosophila* community, in particular David Strutt, Marek Mlodzik and Yohanns Bellaiche, and the Bloomington Stock center for reagents. The Consolidated Groups Program of the Generalitat de Catalunya and DGI and Consolider Grants from the Ministry of Economy and Competitivity of Spain to EMB supported this work. FM was a JAE-CSIC predoctoral fellow.

## AUTHOR CONTRIBUTION

FM performed all experimental procedures; FM and EMB designed the study, analyzed the data and wrote the paper.

## SUPPLEMENTARY FIGURES

**Figure S1.**
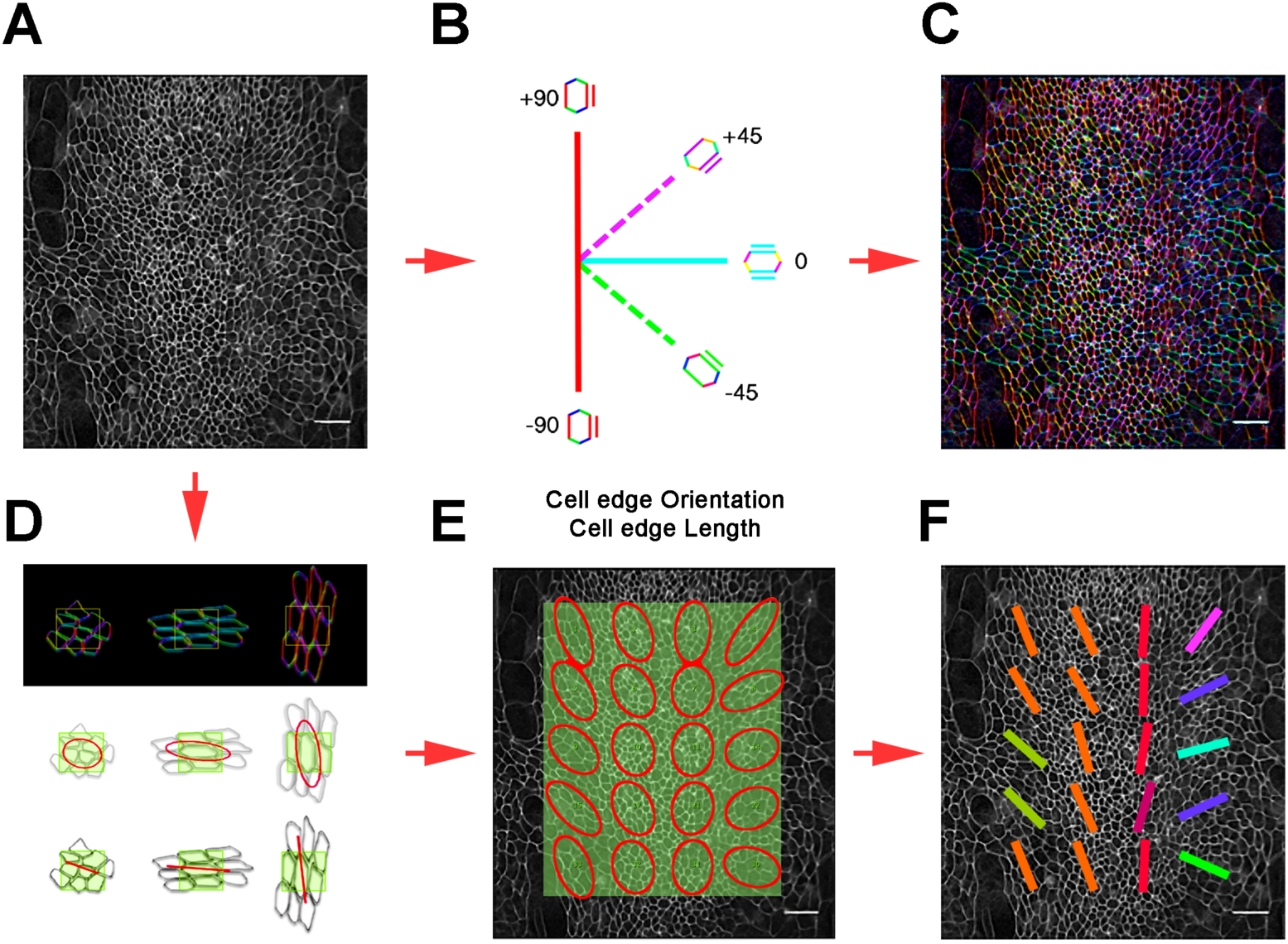
Determination of cell edges alignment orientation and coherency. **A**) Images (snapshots) of abdominal cells outlined with ZCL2207 were analyzed with Orientation J (Fiji) as indicated in the Experimental Procedures. **B**) The coordinate system defined and color-coded with Orientation J was linked to the A/P boundary (± 90° - red) or orthogonal to it (0° - cyan). Orientations tilting clockwise from +90° to 0° have positive values (+45° - magenta). Orientations tilting counterclockwise from - 90° to 0° have negative values (−45° - green). **C**) Full color-coded representation of the orientation of cell edges. **D**) Full images were employed to extract local averaged values for the orientation and coherency of cell edges and specific ROIs were selected for averaged measurements. **E**) The software renders the numerical values for orientation angles and coherency and fit them into ellipses positioned on the ROIs coordinates. F) Orientation Angles values were color coded with the orientation LUT described above and represented on the topographical space of the analyzed areas.

**Figure S2.**
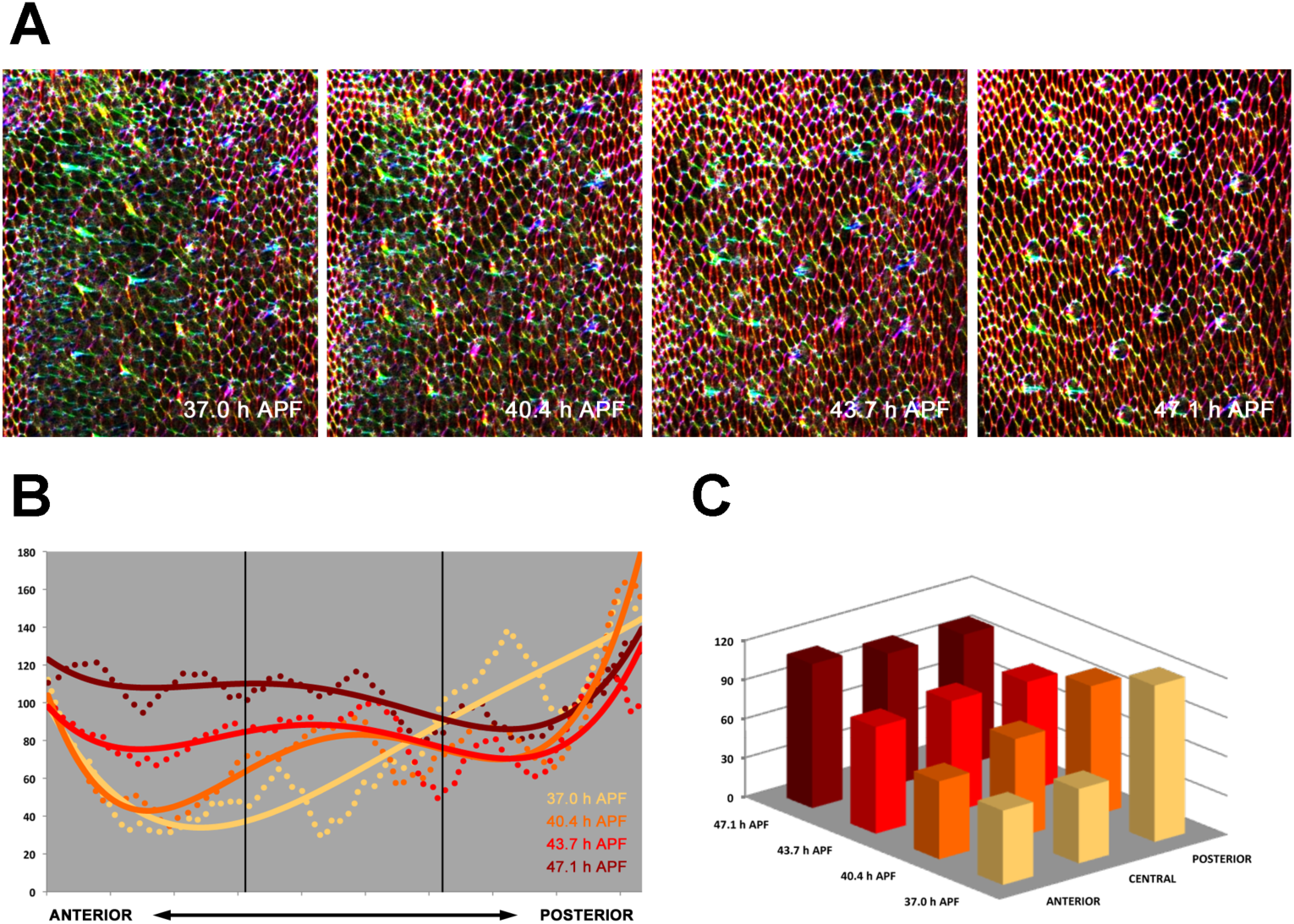
Topographical and temporal axial orientation of cell alignment through the epithelial landscape. **A)** 2D ROIs across the anterior compartment of the dorsal anterior histoblast nest selected at 4 different time points (every 3.35 hours) throughout the long-range orientation of the epithelium. Anterior is to the left and dorsal is up. Scale bar is 22 μm. **B)** Normalized values for cells orientation plotted along the A/P axis (see Experimental Procedures) at different time points (color coded from light to dark browness as stated in the image). Dashed lines represent the raw data and continuous lines their 4th order polynomial regression. An orientation wave generating at the posterior edge of the compartment was readily observed. Oscillatory alterations of intensity level were caused by the periodical positioning of sensory organs, which locally affect cells orientations. **C)** Averages of normalized orientation values at anterior, central and posterior domains in color-coded 3D histograms. As long-range remodeling proceeds, a directed axial progression towards uniformity in orientation develops. All cells sequentially align parallel to the A/P axis from posterior to anterior, creating a uniform landscape.

**Figure S3.**
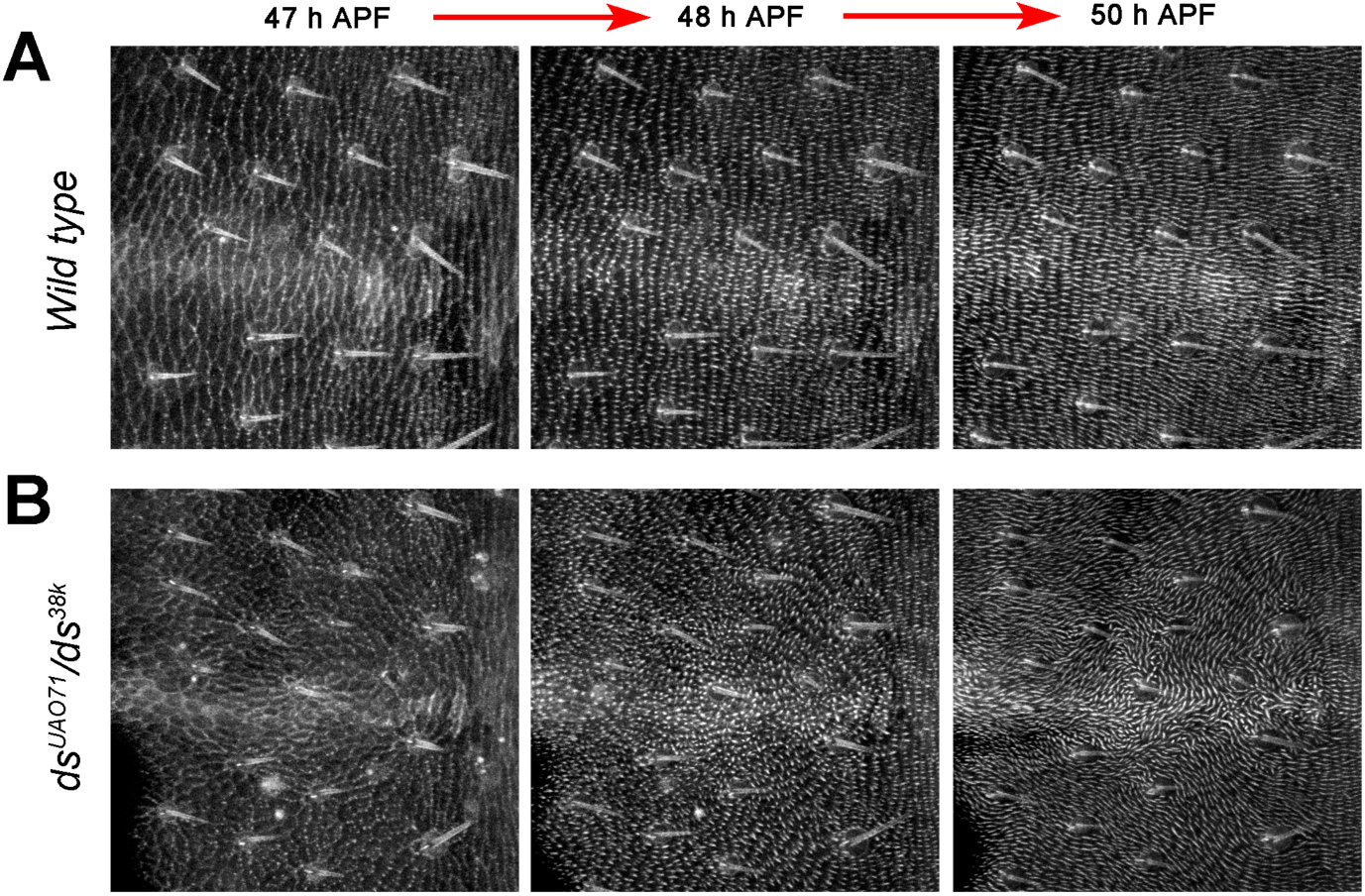
Trichomes patterns in *wt* and *ds* pupae. **A)** Trichomes eversión and growth in *wt* pupal abdomens between 45 and 50 h APF (dorsal view). Actin (moesin∷GFP) accumulates at the posterior side of each cell resolving into up to 5 posteriorly pointing trichomes per cell. A uniform pattern of oriented trichomes is observed from 50 h APF. **B**) Trichomes eversion and orientation in *ds (ds^UAOD71^/ds^38K^)* mutant pupal abdomen between 45 and 50 h APF. Trichomes (moesin∷GFP), as in *wt* animals, initiate at the apical cell periphery of most of the cells. Despite the failure in long-range cell alignment orientation, actin accumulation proceeds parallel to the cells long axis in all cases. Trichomes outgrowth follows the same temporal dynamics as in *wt* but they do not uniformly point posterior. Consequently, a swirling pattern of oriented hairs is observed from 50 h APF. Scale bar is 5 μm. Anterior is to the left.

**Figure S4.**
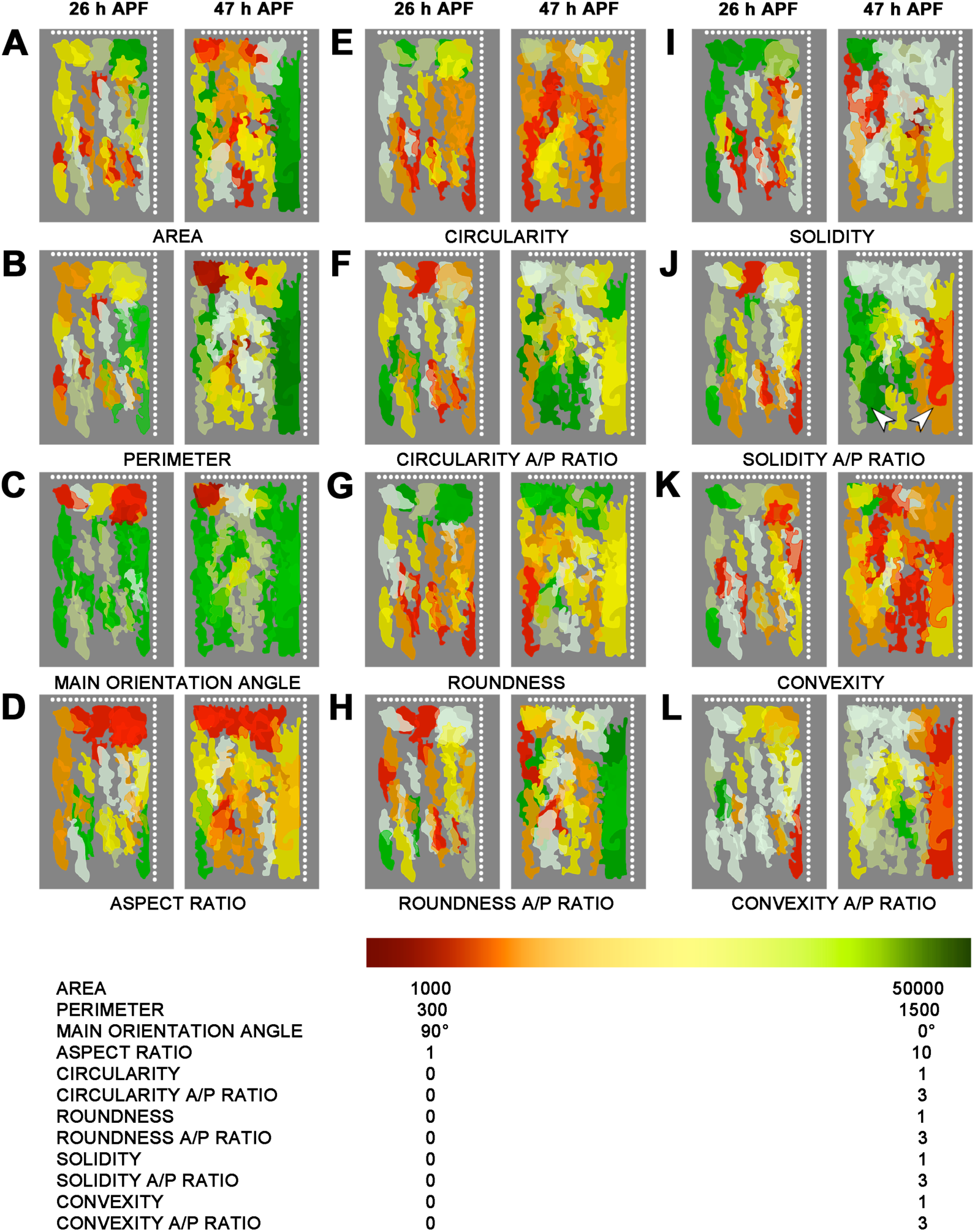
Clonal geometrical and shape parameters. Geometrical and shape parameters were determined for *wt* clones *wt* clones analyzed at 26 and 47 h APF. Cumulative topographic plots displaying the positions and shapes of individual clones in an idealized A compartment (dorsal and posterior edges are represented by dotted white lines). Individual clones were color-coded according to a graded Green-Red LUTs. **A)** Area. **B**) Perimeter. **C)** Main orientation angle. **D)** Aspect Ratio. **E)** Circularity. **F)** Circularity A/P Ratio. **G)** Roundness. **H)** Roundness A/P Ratio. **I)** Solidity. **J)** Solidity A/P Ratio. **K)** Convexity. **L)** Convexity A/P Ratio. The bias in Solidity A/P Ratio at different positions in the compartment is highlighted (arrowheads). The different maximum and minimum values for each analyzed parameter are stated at the bottom legend (see Experimental Procedures). For some parameters (Circularity, Roundness, Solidity and Convexity), values were calculated for the anterior and posterior bisected halves of each clone and their ratio was color-coded for each clone as above.

## SUPPLEMENTARY MOVIES LEGENDS

**Movie S1: Tissue-wide changes in cell orientation pattern during the early epithelial expansion**

Confocal time-lapse of an Atpα∷GFP *Drosophila* pupae depicting cell junctions of histoblasts and LECs. The video run from 16 to 26 h APF and displays a dorso lateral view of the expansion of the anterior and posterior dorsal histoblast nests of a third thoracic segment. Color-coded cell outlines on the right show the orientation of the cell interfaces relative to the A/P boundary. Red indicates cell interfaces oriented in parallel to the A/P boundary. Cyan depicts cell interfaces oriented perpendicular to it. Other color-coded angles are indicated. Note over the time course of the movie the increased proportion of A/P oriented interfaces (red) in the vicinity of the A/P boundary. Scale bar is 22 μm. Anterior is to the left and dorsal is up. Time intervals were 300 seconds. See Figs. 1A and S1.

**Movie S2: Tissue-wide changes in cell orientation pattern during the late epithelial remodeling**

Confocal time-lapse of an Atpα∷GFP *Drosophila* pupae depicting cell junctions of histoblasts and LECs. The video run from 38 to 47 h APF and displays a dorsal view of the remodeling of the contralateral dorsal histoblast nests of a third thoracic segment. Color-coded cell outlines on the right show the orientation of the cell interfaces relative to the A/P boundary. Red indicates cell interfaces oriented in parallel to the A/P boundary. Cyan depicts cell interfaces oriented perpendicular to it. Other coded-coded angles are indicated. Note the progression from posterior to anterior of the long-range A/P orientation of the tissue. At the end of remodeling all cells orient in alignment with the A/P boundary. Scale bar is 22 μm. Anterior is to the left and dorsal is up. Time intervals were 300 seconds. See Figs. 1B and S1.

**Movie S3: Tissue-wide changes in cell orientation pattern are perturbed in dachsous pupae during epithelial expansion**

Confocal time-lapse of an Atpα∷GFP *Drosophila ds* pupae depicting cell junctions of histoblasts and LECs. The video run from 16 to 26 h APF and displays the expansion of the anterior and posterior dorsal histoblast nests of a third thoracic segment. Color-coded cell outlines on the right show the orientation of the cell interfaces relative to the A/P boundary. Red indicates cell interfaces oriented in parallel to the A/P boundary. Cyan depicts cell interfaces oriented perpendicular to it. Other coded-coded angles are indicated. Note that just a few cells close to the A/P boundary become oriented by 26 h APF. Scale bar is 22 μm. Anterior is to the left and dorsal is up. Time intervals were 300 seconds. See Fig. 2A for quantifications.

**Movie S4: A uniform tissue-wide pattern of cell orientation fails to emerge in *dachsous* pupae during epithelial remodeling**

Confocal time-lapse of an Atpα∷GFP *Drosophila ds* pupae depicting cell junctions of histoblasts and LECs. The video run from 38 to 47 h APF and displays a dorsal view of the remodeling of the contralateral dorsal histoblast nests of a third thoracic segment. Color-coded cell outlines on the right show the orientation of the cell interfaces relative to the A/P boundary. Red indicates cell interfaces oriented in parallel to the A/P boundary. Cyan depicts cell interfaces oriented perpendicular to it. Other color-coded angles are indicated. Note that just a few cells close to the A/P boundary become oriented by 26 h APF. Scale bar 22 μm. Anterior is to the left and dorsal is up. Time intervals were 300 seconds. See Fig. 2A for quantifications. Note that long-range cell orientation is not achieved at the end of morphogenesis. Scale bar is 22 μm. Anterior is to the left and dorsal is up. Time intervals were 300 seconds.

**Movie S5. The emergence of a uniform pattern of trichome orientation requires uniform pattern of cell orientation**

High magnification time-lapse depicting the ubiquitous expression of a Moesin∷GFP actin reporter between 45 and 50 h APF in an otherwise *wt* pupae. Note the emergence of trichomes perpendicular to the long axis of each cell and their orientation perpendicular to the A/P boundary.

The video displays a dorsal ROIs within an anterior dorsal nest of the third abdominal segment. On the right, the orientation of trichome outgrowth relative to the A/P boundary is color-coded. Red indicates orientation parallel to the A/P boundary, while cyan depicts perpendicular orientation. Other color-coded angles are indicated. Scale bar is 5 μm. Anterior is to the left and dorsal is up. Time intervals were 300 seconds.

**Movie S6. The emergence of a uniform pattern of trichome orientation is disrupted in *ds* pupae**

High magnification video time-lapse depicting the ubiquitous expression of a Moesin∷GFP actin reporter between 45 and 50 h APF in a *ds* pupae. Note that trichome development keeps perpendicular to the cells longer axis but is no longer perpendicular to the A/P boundary.

The video displays a dorsal ROIs within an anterior dorsal nest of the third abdominal segment. On the right, the orientation of trichome outgrowth relative to the A/P boundary is color-coded. Red indicates orientation parallel to the A/P boundary, while cyan depicts perpendicular orientation. Other color-coded angles are indicated. Scale bar is 5 μm. Anterior is to the left and dorsal is up. Time intervals were 300 seconds.

**Movie S7: Expression pattern dynamics of Ds during epithelial morphogenesis**

**A**) Video time-lapse depicting the endogenous expression pattern of Dachsous (Ds∷EGFP) during histoblasts expansion between 16 and 26 h APF in an otherwise *wt* pupae. The video displays the anterior and posterior dorsal histoblast nests of a third thoracic segment. **B)** Video time-lapse depicting the endogenous expression pattern of Dachsous (Ds∷EGFP) during tissue remodeling between 38 and 46 h APF in an otherwise *wt* pupae. The video displays a dorsal view of the contralateral dorsal histoblast nests of the third abdominal segment.

Note that by 16 h APF, Ds∷EGFP is expressed in gradients with highest levels at the A/P boundary. During morphogenesis, Ds gradients get shallower, but sustain their orientation. Scale bar is 22 μm. See also Fig. 4.

**Movie S8: Expression pattern dynamics of *fj* during epithelial morphogenesis**

**A)** Video time-lapse depicting the endogenous expression pattern of *four-jointed (fj-* GFP) during histoblasts expansion between 16 and 26 h APF in an otherwise *wt* pupae. The video displays the anterior and posterior dorsal histoblast nests of a third thoracic segment. **B)** Video time-lapse depicting the endogenous expression pattern of *fj*-GFP during tissue remodeling between 38 and 46 h APF in an otherwise *wt* pupae. The video displays a dorsal view of the contralateral dorsal histoblast nests of the third abdominal segment.

Note that by 16 h APF, *fj^CB04634^* is expressed in opposing gradients in the A and P compartments with highest levels at the segmental borders and a minimum at the A/P boundary. During morphogenesis, these gradients get shallower, but sustain their orientation. Scale bar is 22 μm. See also Fig. 4.

**Movie S9: Expression pattern dynamics of Ft during epithelial morphogenesis**

**A)** Confocal time-lapse depicting the endogenous expression pattern of Fat (Ft∷EGFP) during histoblasts expansion between 16 and 26 h APF in an otherwise *wt* pupae. The video displays the anterior and posterior dorsal histoblast nests of a third thoracic segment. **B)** Video time-lapse depicting the endogenous expression pattern of Fat (Ft∷EGFP) during tissue remodeling between 38 and 46 h APF in an otherwise *wt* pupae. The video displays a dorsal view of the contralateral dorsal histoblast nests of the third abdominal segment.

Note that between 16 and 26 h APF, Ft∷EGFP is not expressed in a gradient. However, at remodeling its expression evolves into gradients opposing those of Ds. During morphogenesis, these gradients get shallower, but sustain their orientation. Scale bar is 22 μm. See also Fig. 4.

## REFERENCES

Aigouy, B., Farhadifar, R., Staple, D.B., Sagner, A., Roper, J.C., Julicher, F., and Eaton, S. (2010). Cell flow reorients the axis of planar polarity in the wing epithelium of Drosophila. Cell 142, 773–786.

Amack, J.D., and Manning, M.L. (2012). Knowing the boundaries: extending the differential adhesion hypothesis in embryonic cell sorting. Science 338, 212–215.

Ambegaonkar, A.A., Pan, G., Mani, M., Feng, Y., and Irvine, K.D. (2012). Propagation of Dachsous-Fat planar cell polarity. Current biology: CB 22, 1302–1308.

Bainbridge, S.P., and Bownes, M. (1981). Staging the metamorphosis of Drosophila melanogaster. Journal of embryology and experimental morphology 66, 57–80.

Beloussov, L.V. (2015). Morphomechanics of Development (Springer).

Bischoff, M., and Cseresnyes, Z. (2009). Cell rearrangements, cell divisions and cell death in a migrating epithelial sheet in the abdomen of Drosophila. Development 136, 2403–2411.

Boote, C., Kamma-Lorger, C.S., Hayes, S., Harris, J., Burghammer, M., Hiller, J., Terrill, N.J., and Meek, K.M. (2011). Quantification of collagen organization in the peripheral human cornea at micron-scale resolution. Biophysical journal 101, 33–42.

Bosveld, F., Bonnet, I., Guirao, B., Tlili, S., Wang, Z., Petitalot, A., Marchand, R., Bardet, P.L., Marcq, P., Graner, F., et al. (2012). Mechanical control of morphogenesis by Fat/Dachsous/Four-jointed planar cell polarity pathway. Science 336, 724–727.

Bourget, J.-M., Auger, F.A., Germain, L., Guillemette, M., and Veres, T. (2013). Alignment of Cells and Extracellular Matrix Within Tissue-Engineered Substitutes (INTECH Open Access Publisher).

Brittle, A., Thomas, C., and Strutt, D. (2012). Planar polarity specification through asymmetric subcellular localization of Fat and Dachsous. Current biology: CB 22, 907–914.

Brittle, A.L., Repiso, A., Casal, J., Lawrence, P.A., and Strutt, D. (2010). Four-jointed modulates growth and planar polarity by reducing the affinity of dachsous for fat. Current biology: CB 20, 803–810.

Campinho, P., Behrndt, M., Ranft, J., Risler, T., Minc, N., and Heisenberg, C.P. (2013). Tension-oriented cell divisions limit anisotropic tissue tension in epithelial spreading during zebrafish epiboly. Nature cell biology 15, 1405–1414.

Carvajal-Gonzalez, J.M., and Mlodzik, M. (2014). Mechanisms of planar cell polarity establishment in Drosophila. F1000prime reports 6, 98.

Casal, J., Lawrence, P.A., and Struhl, G. (2006). Two separate molecular systems, Dachsous/Fat and Starry night/Frizzled, act independently to confer planar cell polarity. Development 133, 4561–4572.

Casal, J., Struhl, G., and Lawrence, P.A. (2002). Developmental compartments and planar polarity in Drosophila. Current biology: CB 12, 1189–1198.

Clark, H.F., Brentrup, D., Schneitz, K., Bieber, A., Goodman, C., and Noll, M. (1995). Dachsous encodes a member of the cadherin superfamily that controls imaginal disc morphogenesis in Drosophila. Genes & Development 9, 1530–1542.

Devenport, D. (2014). The cell biology of planar cell polarity. The Journal of cell biology 207, 171–179.

Fagotto, F. (2014). The cellular basis of tissue separation. Development 141, 3303–3318.

Fagotto, F. (2015). Regulation of cell adhesion and cell sorting at embryonic boundaries. Current topics in developmental biology 112, 19–64.

Fonck, E., Feigl, G.G., Fasel, J., Sage, D., Unser, M., Rufenacht, D.A., and Stergiopulos, N. (2009). Effect of aging on elastin functionality in human cerebral arteries. Stroke; a journal of cerebral circulation 40, 2552–2556.

Foty, R.A., and Steinberg, M.S. (2005). The differential adhesion hypothesis: a direct evaluation. Developmental biology 278, 255–263.

Gao, B. (2012). Wnt regulation of planar cell polarity (PCP). Current topics in developmental biology 101, 263–295.

Garcia-Bellido, A., and Merriam, J.R. (1971). Clonal parameters of tergite development in Drosophila. Developmental biology 26, 264–276.

Garoia, F., Guerra, D., Pezzoli, M.C., Lopez-Varea, A., Cavicchi, S., and Garcia-Bellido, A. (2000). Cell behaviour of Drosophila fat cadherin mutations in wing development. Mech Dev 94, 95–109.

Golic, K.G., and Lindquist, S. (1989). The FLP recombinase of yeast catalyzes site-specific recombination in the Drosophila genome. Cell 59, 499–509.

Hale, R., Brittle, A.L., Fisher, K.H., Monk, N.A., and Strutt, D. (2015). Cellular interpretation of the long-range gradient of Four-jointed activity in the Drosophila wing. eLife 4.

Hale, R., and Strutt, D. (2015). Conservation of Planar Polarity Pathway Function Across the Animal Kingdom. Annual review of genetics 49, 529–551.

Hammer, Ø., Harper, D., and Ryan, P. (2001). PAST-Palaeontological statistics. www.uves/~pardomv/pe/2001_1/past/pastprog/past.pdf, acessado em 25, 2009.

Hayashi, T., and Carthew, R.W. (2004). Surface mechanics mediate pattern formation in the developing retina. Nature 431, 647–652.

Heisenberg, C.P., and Bellaiche, Y. (2013). Forces in tissue morphogenesis and patterning. Cell 153, 948–962.

Holzapfel, G.A., Sommer, G., Gasser, C.T., and Regitnig, P. (2005). Determination of layer-specific mechanical properties of human coronary arteries with nonatherosclerotic intimal thickening and related constitutive modeling. American Journal of Physiology-Heart and Circulatory Physiology 289, H2048–H2058.

Hopyan, S., Sharpe, J., and Yang, Y. (2011). Budding behaviors: Growth of the limb as a model of morphogenesis. Developmental dynamics: an official publication of the American Association of Anatomists 240, 1054–1062.

Ishikawa, H.O., Takeuchi, H., Haltiwanger, R.S., and Irvine, K.D. (2008). Four-jointed is a Golgi kinase that phosphorylates a subset of cadherin domains. Science 321, 401–404.

Kafer, J., Hayashi, T., Maree, A.F., Carthew, R.W., and Graner, F. (2007). Cell adhesion and cortex contractility determine cell patterning in the Drosophila retina. Proceedings of the National Academy of Sciences of the United States of America 104, 18549–18554.

Keller, R. (2006). Mechanisms of elongation in embryogenesis. Development 133, 2291–2302.

Kornberg, T. (1981). Compartments in the abdomen of Drosophila and the role of the engrailed locus. Developmental biology 86, 363–372.

Lawrence, P.A., and Casal, J. (2013). The mechanisms of planar cell polarity, growth and the Hippo pathway: some known unknowns. Developmental biology 377, 1–8.

Lawrence, P.A., Casal, J., and Struhl, G. (1999). The hedgehog morphogen and gradients of cell affinity in the abdomen of Drosophila. Development 126, 2441–2449.

Lawrence, P.A., Casal, J., and Struhl, G. (2002). Towards a model of the organisation of planar polarity and pattern in the Drosophila abdomen. Development 129, 2749–2760.

Lecuit, T., and Lenne, P.F. (2007). Cell surface mechanics and the control of cell shape, tissue patterns and morphogenesis. Nature reviews Molecular cell biology 8, 633–644.

Lecuit, T., Lenne, P.F., and Munro, E. (2011). Force generation, transmission, and integration during cell and tissue morphogenesis. Annual review of cell and developmental biology 27, 157–184.

Legoff, L., Rouault, H., and Lecuit, T. (2013). A global pattern of mechanical stress polarizes cell divisions and cell shape in the growing Drosophila wing disc. Development 140, 4051–4059.

Lynch, H.A., Johannessen, W., Wu, J.P., Jawa, A., and Elliott, D.M. (2003). Effect of fiber orientation and strain rate on the nonlinear uniaxial tensile material properties of tendon. Journal of biomechanical engineering 125, 726–731.

Ma, D., Yang, C.H., McNeill, H., Simon, M.A., and Axelrod, J.D. (2003). Fidelity in planar cell polarity signalling. Nature 421, 543–547.

Madhavan, M.M., and Madhavan, K. (1980). Morphogenesis of the epidermis of adult abdomen of Drosophila. Journal of embryology and experimental morphology 60, 1–31.

Madhavan, M.M., and Schneiderman, H.A. (1977). Histological analysis of the dynamics of growth of imaginal discs and histoblast nests during the larval development ofDrosophila melanogaster. Wilhelm Roux’s archives of developmental biology 183, 269–305.

Mao, Y., Rauskolb, C., Cho, E., Hu, W.L., Hayter, H., Minihan, G., Katz, F.N., and Irvine, K.D. (2006). Dachs: an unconventional myosin that functions downstream of Fat to regulate growth, affinity and gene expression in Drosophila. Development 133, 2539–2551.

Mao, Y., Tournier, A.L., Bates, P.A., Gale, J.E., Tapon, N., and Thompson, B.J. (2011). Planar polarization of the atypical myosin Dachs orients cell divisions in Drosophila. Genes Dev 25, 131–136.

Mao, Y., Tournier, A.L., Hoppe, A., Kester, L., Thompson, B.J., and Tapon, N. (2013). Differential proliferation rates generate patterns of mechanical tension that orient tissue growth. The EMBO journal 32, 2790–2803.

Matakatsu, H., and Blair, S.S. (2004). Interactions between Fat and Dachsous and the regulation of planar cell polarity in the Drosophila wing. Development 131, 3785–3794.

Matakatsu, H., and Blair, S.S. (2006). Separating the adhesive and signaling functions of the Fat and Dachsous protocadherins. Development 133, 2315–2324.

Matis, M., and Axelrod, J.D. (2013). Regulation of PCP by the Fat signaling pathway. Genes Dev 27, 2207–2220.

Nardi, J.B., and Kafatos, F.C. (1976). Polarity and gradients in lepidopteran wing epidermis. II. The differential adhesiveness model: gradient of a non-diffusible cell surface parameter. Journal of embryology and experimental morphology 36, 489–512.

Ninov, N., Chiarelli, D.A., and Martin-Blanco, E. (2007). Extrinsic and intrinsic mechanisms directing epithelial cell sheet replacement during Drosophila metamorphosis. Development 134, 367–379.

Ninov, N., and Martin-Blanco, E. (2007). Live imaging of epidermal morphogenesis during the development of the adult abdominal epidermis of Drosophila. Nature protocols 2, 3074–3080.

Nubler-Jung, K. (1987). Tissue polarity in an insect segment: denticle patterns resemble spontaneously forming fibroblast patterns. Development 100, 171–177.

Nubler-Jung, K., and Mardini, B. (1990). Insect epidermis: polarity patterns after grafting result from divergent cell adhesions between host and graft tissue. Development 110, 1071–1079.

Pilot, F., and Lecuit, T. (2005). Compartmentalized morphogenesis in epithelia: from cell to tissue shape. Developmental dynamics: an official publication of the American Association of Anatomists 232, 685–694.

Rauzi, M., Verant, P., Lecuit, T., and Lenne, P.F. (2008). Nature and anisotropy of cortical forces orienting Drosophila tissue morphogenesis. Nature cell biology 10, 1401–1410.

Rezakhaniha, R., Agianniotis, A., Schrauwen, J.T., Griffa, A., Sage, D., Bouten, C.V., van de Vosse, F.N., Unser, M., and Stergiopulos, N. (2012). Experimental investigation of collagen waviness and orientation in the arterial adventitia using confocal laser scanning microscopy. Biomechanics and modeling in mechanobiology 11, 461–473.

Rogulja, D., Rauskolb, C., and Irvine, K.D. (2008). Morphogen control of wing growth through the Fat signaling pathway. Developmental cell 15, 309–321.

Roseland, C.R., and Schneiderman, H.A. (1979). Regulation and metamorphosis of the abdominal histoblasts ofDrosophila melanogaster. Wilhelm Roux’s archives of developmental biology 186, 235–265.

Santamaria, P., and Garcia-Bellido, A. (1972). Localization and growth pattern of the tergite Anlage of Drosophila. Journal of embryology and experimental morphology 28, 397–417.

Sharma, P., and McNeill, H. (2013a). Fat and Dachsous cadherins. Progress in molecular biology and translational science 116, 215–235.

Sharma, P., and McNeill, H. (2013b). Regulation of long-range planar cell polarity by Fat-Dachsous signaling. Development 140, 3869–3881.

Simon, M.A. (2004). Planar cell polarity in the Drosophila eye is directed by graded Four-jointed and Dachsous expression. Development 131, 6175–6184.

Simon, M.A., Xu, A., Ishikawa, H.O., and Irvine, K.D. (2010). Modulation of fat:dachsous binding by the cadherin domain kinase four-jointed. Current biology: CB 20, 811–817.

Strutt, D. (2009). Gradients and the specification of planar polarity in the insect cuticle. Cold Spring Harbor perspectives in biology 1, a000489.

Strutt, H., and Strutt, D. (2002). Nonautonomous planar polarity patterning in Drosophila: dishevelled-independent functions of frizzled. Developmental cell 3, 851–863.

Sugimura, K., and Ishihara, S. (2013). The mechanical anisotropy in a tissue promotes ordering in hexagonal cell packing. Development 140, 4091–4101.

Suzanne, M., Petzoldt, A.G., Speder, P., Coutelis, J.B., Steller, H., and Noselli, S. (2010). Coupling of apoptosis and L/R patterning controls stepwise organ looping. Current biology: CB 20, 1773–1778.

Thomas, C., and Strutt, D. (2012). The roles of the cadherins Fat and Dachsous in planar polarity specification in Drosophila. Developmental dynamics: an official publication of the American Association of Anatomists 241, 27–39.

Umetsu, D., Aigouy, B., Aliee, M., Sui, L., Eaton, S., Julicher, F., and Dahmann, C. (2014). Local increases in mechanical tension shape compartment boundaries by biasing cell intercalations. Current biology: CB 24, 1798–1805.

Xu, T., and Rubin, G.M. (1993). Analysis of genetic mosaics in developing and adult Drosophila tissues. Development 117, 1223–1237.

Yang, C.H., Axelrod, J.D., and Simon, M.A. (2002). Regulation of Frizzled by fat-like cadherins during planar polarity signaling in the Drosophila compound eye. Cell 108, 675–688.

Zeidler, M.P., Perrimon, N., and Strutt, D.I. (1999). The four-jointed gene is required in the Drosophila eye for ommatidial polarity specification. Current biology: CB 9, 1363–1372.

Zeidler, M.P., Perrimon, N., and Strutt, D.I. (2000). Multiple roles for four-jointed in planar polarity and limb patterning. Developmental biology 228, 181–196.

Zhu, H. (2009). Is anisotropic propagation of polarized molecular distribution the common mechanism of swirling patterns of planar cell polarization? Journal of theoretical biology 256, 315–325.

